# Photoreceptor–induced sinapate synthesis contributes to photoprotection in Arabidopsis

**DOI:** 10.1101/2024.02.26.582123

**Authors:** Manuela Leonardelli, Nicolas Tissot, Roman Podolec, Florence Ares-Orpel, Gaétan Glauser, Roman Ulm, Emilie Demarsy

**Affiliations:** Department of Plant Sciences, Section of Biology, Faculty of Sciences, University of Geneva, CH-1211 Geneva 4, Switzerland; Institute of Genetics and Genomics of Geneva (iGE3), University of Geneva, Geneva, Switzerland; Neuchâtel Platform of Analytical Chemistry, University of Neuchâtel, CH-2000 Neuchâtel, Switzerland

## Abstract

Plants must balance light capture for photosynthesis with protection from potentially harmful ultraviolet radiation (UV). Photoprotection is mediated by concerted action of photoreceptors, but the underlying molecular mechanisms are not fully understood. In this study, we provide evidence that UV RESISTANCE LOCUS 8 (UVR8) UV-B-, phytochrome red-, and cryptochrome blue-light photoreceptors converge on the induction of *FERULIC ACID 5-HYDROXYLASE 1* (*FAH1*) that encodes a key enzyme in the phenylpropanoid biosynthesis pathway, leading to the accumulation of UV-absorbing sinapate esters. *FAH1* induction depends on the bZIP transcription factors ELONGATED HYPOCOTYL 5 (HY5) and HY5-HOMOLOG (HYH) that function downstream of all three photoreceptors. Noticeably, mutants with hyperactive UVR8 signalling rescue *fah1* UV sensitivity. Targeted metabolite profiling suggests that this phenotypic rescue is due to the accumulation of UV-absorbing metabolites derived from precursors of sinapate synthesis, namely coumaroyl-glucose and feruloyl-glucose. Our genetic dissection of the phenylpropanoid pathway combined with metabolomic and physiological analyses show that both sinapate esters and flavonoids contribute to photoprotection with sinapates playing a major role for UV screening. Our findings indicate that photoreceptor-mediated regulation of *FAH1* and subsequent accumulation of sinapate “sunscreen” compounds is a key protective mechanism to mitigate damage, preserving photosynthetic performance, and ensuring plant survival under UV.

## INTRODUCTION

Light fuels photosynthesis and affects plant growth, development, and metabolism throughout their life cycle. To optimize growth and development according to ambient light conditions, plants employ a set of photoreceptors, including the red/far-red–sensory phytochromes (phyA– phyE); the blue light–perceiving cryptochromes (cry1, cry2); and the ultraviolet-B (UV-B)/short wavelength UV-A radiation photoreceptor UV RESISTANCE LOCUS 8 (UVR8) (Galvao and Fankhauser, 2015; Allorent and Petroutsos, 2017; Demarsy et al., 2018; Rai et al., 2020; Podolec et al., 2021a). Upon light exposure, phytochrome, cryptochrome, and UVR8 photoreceptors inhibit the E3 ubiquitin ligase CONSTITUTIVELY PHOTOMORPHOGENIC 1 (COP1), thereby allowing stabilization of key transcription factors involved in photomorphogenic responses such as the basic leucine zipper (bZIP) transcription factors ELONGATED HYPOCOTYL 5 (HY5) and HY5 HOMOLOG (HYH) (Osterlund et al., 2000; Holm et al., 2002; Favory et al., 2009; Binkert et al., 2014; Yin et al., 2015; Hoecker, 2017; Podolec and Ulm, 2018; Lau et al., 2019; Ponnu et al., 2019).

UVR8 exists as a homodimer in its ground state (Favory et al., 2009; Rizzini et al., 2011). Upon UV-B photon absorption, UVR8 monomerizes and accumulates in the nucleus in a COP1-dependent manner (Kaiserli and Jenkins, 2007; Rizzini et al., 2011; Yin et al., 2016; Fang et al., 2022). The two WD40-repeat proteins REPRESSOR OF UV-B PHOTOMORPHOGENESIS 1 (RUP1) and RUP2 act as negative feedback regulators of the UVR8 photocycle by facilitating UVR8 re-dimerization and thereby repressing UV-B signalling (Gruber et al., 2010; Heijde and Ulm, 2013; Wang et al., 2023). In agreement, *rup1 rup2* mutants show enhanced UV-B responsiveness and acclimation (Gruber et al., 2010). Interestingly, the *uvr8-17D* mutant, which expresses the constitutively monomeric UVR8^G101S^ photoreceptor variant, phenotypically resembles *rup1 rup2*, reinforcing the importance of UVR8 redimerization to optimally balance UV-B-induced photomorphogenesis with plant growth and development (Podolec et al., 2021b).

Despite its necessity for photosynthesis, light can also constitute an environmental stressor for the photosynthetic machinery when photosynthetic capacity is overwhelmed, as well as due to its intrinsic, potentially damaging UV-B component (Jansen et al., 1998; Li et al., 2009; Demarsy et al., 2018). UV-B induced damage to photosystem II (PSII) occurs primarily in the water-oxidizing manganese cluster of the oxygen evolving complex, and at the D1 and D2 proteins (Takahashi et al., 2010; Takahashi and Badger, 2011; Demarsy et al., 2018). Photoreceptors can promote photoprotection, a set of mechanisms through which plants alleviate the negative effects of light on cell integrity and particularly on the photosynthetic machinery at PSII (Jung and Niyogi, 2008; Allorent and Petroutsos, 2017; Demarsy et al., 2018). Evidence in Arabidopsis and *Chlamydomonas reinhardtii* suggests that UVR8 controls an acclimatory response that maintains optimal photosynthetic efficiency under elevated UV-B (Davey et al., 2012; Tilbrook et al., 2016). Recent studies indicate that UVR8 acts in concert with other photoreceptors, mainly cry1, to ensure survival under UV containing sunlight (Morales et al., 2013; Rai et al., 2019; Rai et al., 2020; Tissot and Ulm, 2020; Stockenhuber et al., 2024), but the underlying mechanism has remained enigmatic.

UVR8 signalling leads to transcriptional regulation of genes involved in photomorphogenic responses, including those resulting in UV acclimation and tolerance (Kliebenstein et al., 2002; Brown et al., 2005; Oravecz et al., 2006; Favory et al., 2009; Robson et al., 2019; Podolec et al., 2021a; Rai et al., 2021). Phenolic “sunscreen” metabolites that filter out UV and prevent damage of underlying tissues are important for UV tolerance of land plants (Caldwell et al., 1983; Landry et al., 1995; Sheahan, 1996; Kliebenstein et al., 2002; Stracke et al., 2010; Neugart et al., 2021; Shi and Liu, 2021; Gonzalez Moreno et al., 2022; Procko et al., 2022). “Sunscreen” metabolites in flowering plants mainly include flavonol glycosides and sinapate esters (Tevini et al., 1991; Sheahan, 1996; Burchard et al., 2000), with biosynthesis of flavonol glycosides in particular being known to be UV-B inducible (Li et al., 1993; Landry et al., 1995; Shirley et al., 1995; Stracke et al., 2010; Clayton et al., 2018; Neugart et al., 2019; Nichelmann and Pescheck, 2021; Shi and Liu, 2021). Flavonols are widely considered as central to UV-B-induced UV photoprotection (e.g. Wellmann, 1975; Caldwell et al., 1983; Ryan et al., 2001; Winkel-Shirley, 2002; Agati and Tattini, 2010; Vogt, 2010; Kusano et al., 2011; Jenkins, 2014; Nichelmann and Pescheck, 2021; Podolec et al., 2021a; Shi and Liu, 2021). By contrast, sinapate esters are generally thought to be constitutively present and provide basal UV tolerance (Hectors et al., 2007; Gotz et al., 2010; Nichelmann and Pescheck, 2021), although UVR8–mediated sinapate accumulation has been reported to contribute to pathogen resistance (Demkura and Ballare, 2012).

Sinapate esters belong to the group of hydroxycinnamic acids (HCAs), a widely occurring class of phenylpropanoids in the plant kingdom. The three major sinapate esters in Arabidopsis are sinapoyl glucose, sinapoyl malate, and sinapoyl choline (Fraser and Chapple, 2011). A key enzyme essential for the biosynthesis of sinapate, the precursor of sinapate esters, is ferulic acid-5-hydroxylase (F5H), a cytochrome P450 (CYP84A1) encoded by the *FAH1* locus in Arabidopsis (Lorenzen et al., 1996; Ruegger et al., 1999). *fah1* mutants are hypersensitive to UV stress, further supporting that sinapate esters play a major role in basal, constitutive UV protection (Landry et al., 1995).

Here, we show that UVR8-mediated UV acclimation alleviates UV-B-induced damage to the photosynthetic machinery by mitigating the damage at PSII. This photoprotection mechanism particularly relies on induced F5H activity and subsequent accumulation of sinapate esters that reduce the penetration of UV into photosynthetic tissue. We further show that hyperactive UV-B signalling in *uvr8-17D* and *rup1 rup2* can suppress the UV-stress hypersensitivity phenotype of *fah1*, likely due to enhanced accumulation of UV-absorbing metabolites derived from sinapate precursors, which arise due to an impaired sinapate biosynthesis pathway in *fah1*. Finally, our data suggest that phytochrome, cryptochrome, and UVR8 photoreceptors orchestrate photoprotection against UV via combined induction of *FAH1* and sinapate ester accumulation.

## RESULTS

### UVR8-mediated UV-B acclimation involves photoprotection of the photosynthetic machinery

UVR8 activation promotes tolerance to UV stress (Kliebenstein et al., 2002; Favory et al., 2009; Podolec et al., 2021a). To quantitatively assess the effect of UVR8 activation and UVR8-mediated acclimation on UV stress–induced photoinhibition, we measured chlorophyll fluorescence in wild type and UVR8 signalling mutants exposed to different UV-B treatments, and determined Fv/Fm ratios representing the maximum quantum yield of PSII (Baker, 2008; Murchie and Lawson, 2013). First, 7-d-old light grown seedlings of Col wild type, *rup1 rup2*, and *uvr8* plants were exposed (acclimated, +/-) or not (-/-) to UV-B for 3 days with supplementary narrowband UV-B (0.08 mW cm^-2^; Figure 1A). At day 10, acclimated and non-acclimated plants were exposed to 2h of broadband UV stress irradiation (non-acclimated and stressed -/+; acclimated and stressed +/+) (2.2 mW cm^-2^). Col wild type, *rup1 rup2*, and *uvr8* grown in white light (-/-) all displayed an optimal Fv/Fm value of about 0.83 (Figure 1B, *upper left*, Figure 1C, *black bars*), characteristic of unstressed Arabidopsis plants (Baker, 2008; Murchie and Lawson, 2013). UV stress on non-acclimated plants (-/+) caused a large decrease of Fv/Fm to about 0.42 in all three genotypes, indicating photoinhibition due to severe damage to PSII (Figure 1B, C), further supporting that UVR8 signalling does not contribute to acute UV stress responses (Gonzalez Besteiro et al., 2011). UV-B acclimation alone (+/-) did not significantly affect Fv/Fm in wild type Col and *rup1 rup2*, but led to a decrease in Fv/Fm in the *uvr8* mutant (Figure 1B, C), indicating that already acclimatory levels of UV-B can damage PSII in the absence of the UVR8 photoreceptor, further demonstrating the crucial role of UVR8 for UV-B acclimation and UV stress tolerance. Moreover, results of UV-stress treatment after acclimation (+/+) directly supported a UVR8-dependent photoprotective effect of UV-B acclimation: in Col wild type, acclimation reduced the extent of photoinhibition by the UV-stress treatment, and this photoprotection was further enhanced in *rup1 rup2*, but was absent in *uvr8* (Figure 1B, C; comparing +/+ with -/+). Whereas *hyh* acclimated like wild type, *hy5* and *hy5 hyh* showed reduced photoprotection, with *hy5 hyh* exhibiting UV-stress effects comparable to *uvr8* (Supplemental Figure S1A). By contrast, *uvr8-17D*, expressing UVR8^G101S^ with enhanced UVR8 activity, showed enhanced UV photoprotection (Supplemental Figure S1B) like that seen for *rup1 rup2* (Figure 1B, C). Interestingly, although enhanced UV photoprotection after acclimation was also detectable in *uvr8-17D hy5* and *uvr8-17D hyh* (Supplemental Figure S1B), no acclimatory effect could be detected in *uvr8-17D hy5 hyh* (Supplemental Figure S1B), supporting that HY5 and HYH are key transcription factors mediating UVR8-dependent UV photoprotection.

**Figure 1.**
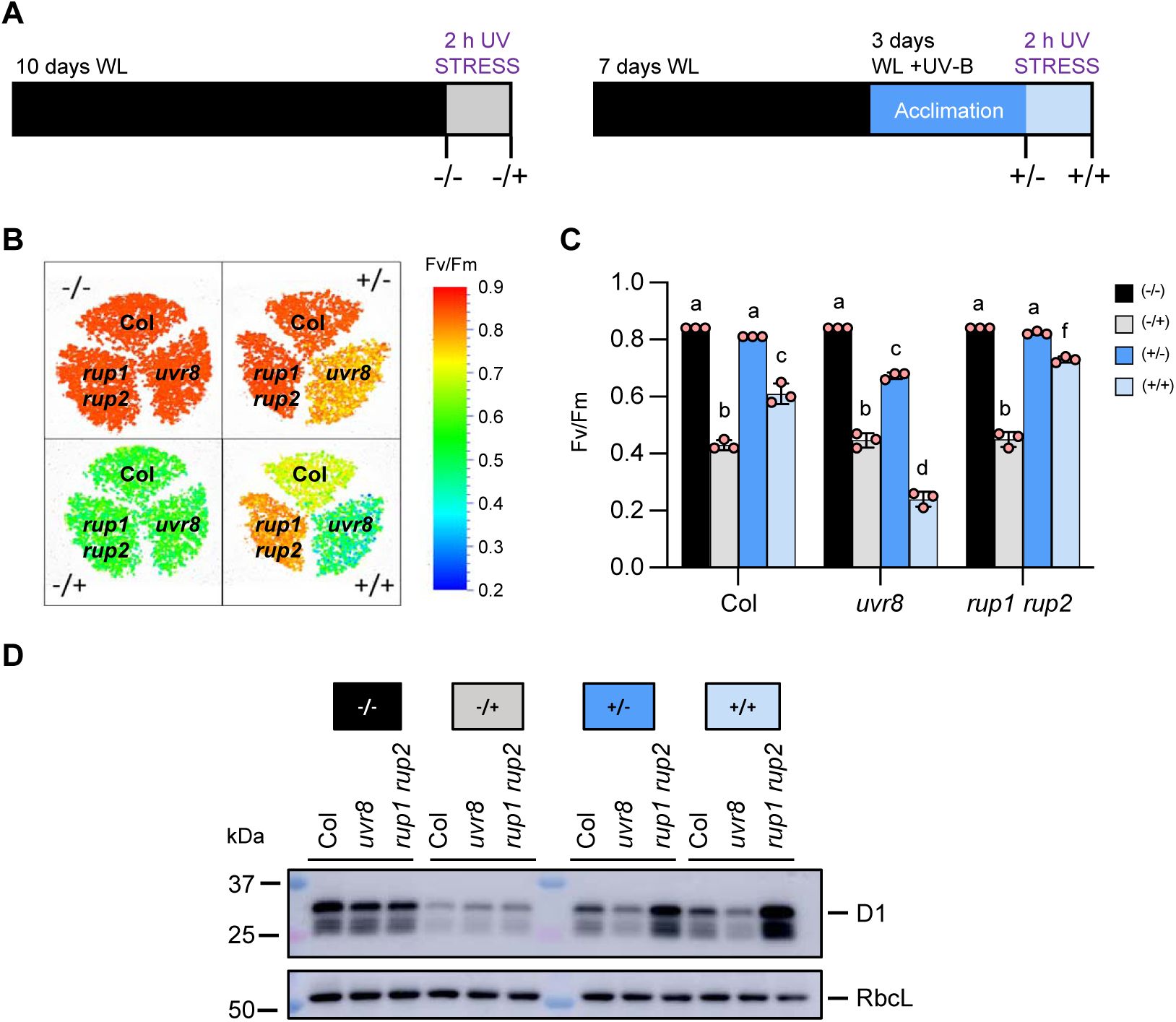
UVR8-dependent UV-B acclimation enhances photoprotection. **(A)** Scheme for UV-B treatment and Fv/Fm sampling. WL (*black bar*): white light (20 µmol m^-2^ s^-1^); WL+UV-B (*blue bar*; Acclimation): WL with supplemental narrowband UV-B (0.08 mW cm^-2^); UV-B STRESS (*grey bar* and *light blue bar*): broadband UV stress (2.2 mW cm^-2^). **(B)** False-color image representing Fv/Fm values of 10-d-old wild type (Col), *uvr8-6* (*uvr8*) and *rup1 rup2* seedlings treated according to the scheme in panel 1A. **(C)** Fv/Fm measurements of 10-d-old wild type (Col), *uvr8-6* (*uvr8*), and *rup1 rup2* seedlings. Values of independent experiments and means ± SD are shown (*n* = 3). Shared letters indicate no statistically significant difference between the means (P > 0.05), as determined by two-way ANOVAs followed by Tukey’s test for multiple comparisons. **(D)** Immunoblot analysis of D1 and RbcL (Rubisco large subunit; loading control) levels in 10-d-old wild type (Col), *uvr8-6*, and *rup1 rup2* seedlings grown and treated as indicated. **(A-D)** -/-, not acclimated, not stressed; -/+, not acclimated, stressed; +/-, acclimated, not stressed; +/+, acclimated, stressed.

To further characterize the effect of UV-B on PSII, we analysed the levels of D1, a PSII core protein and major target of UV-B damage (Booij-James et al., 2000; Takahashi and Badger, 2011; Davey et al., 2012; Tilbrook et al., 2016). UV stress strongly reduced D1 levels across each of wild type Col, *uvr8*, and *rup1 rup2* (-/+ vs. -/-, Figure 1D). Acclimatory UV-B (+/-) reduced D1 levels slightly in Col wild type and more severely in *uvr8*, whereas no reduction was apparent in *rup1 rup2* (Figure 1D), further suggesting that UVR8 is required for photoprotection of the photosynthetic machinery even under acclimatory doses of UV-B. In addition, and in agreement with the Fv/Fm data, UV-B acclimation mitigated the UV-stress effect on D1 levels in Col wild type and *rup1 rup2*, but not in *uvr8* where D1 levels were already low under acclimatory UV-B conditions (+/-and +/+, Figure 1D). We have thus established experimental conditions to evaluate UV acclimation and UV stress tolerance efficiently and quantitatively using Fv/Fm measurements and, together, our data show that UVR8-mediated acclimation results in photoprotection, preventing PSII damage and D1 degradation, thus maintaining photosynthetic performance under UV.

### UVR8 signalling induces *FAH1* expression and accumulation of sinapate esters that underlies photoprotective acclimation

To better understand the underlying mechanism of UVR8-mediated photoprotection, we investigated the contribution of inducible phenylpropanoid “sunscreen” metabolites. UVR8 is well known to promote the accumulation of flavonol glycosides and anthocyanins by upregulating the expression of genes encoding key biosynthetic enzymes, including chalcone synthase (CHS) that catalyses the first committed step in the flavonoid biosynthesis pathway (Kliebenstein et al., 2002; Brown et al., 2005; Favory et al., 2009; Stracke et al., 2010). Sinapate esters, on the other hand, are so far generally considered to provide constitutive UV protection (Gotz et al., 2010; Rai et al., 2019; Nichelmann and Pescheck, 2021). Interestingly, genome-wide expression analyses indicated that *FAH1*, encoding a key enzyme in sinapate biosynthesis and its derivatives sinapate esters, is induced upon UVR8 activation (Favory et al., 2009; Tavridou et al., 2020a). We further analysed *FAH1* expression in 7-d-old seedlings exposed to 3h and 6h narrowband UV-B. *FAH1* expression indeed increased upon UV-B exposure in wild type and to a reduced extent in *hy5* and *hyh* single mutants, whereas expression induction was abolished in *uvr8* and *hy5 hyh* (Figure 2A). *rup1 rup2* and *uvr8-17D* showed slightly elevated and sustained upregulation of *FAH1* mRNA levels (Figure 2A), which was dependent on HY5 and HYH as indicated by an abolished response in *uvr8-17D hy5 hyh* (Supplemental Figure S2A). Next to HY5 and HYH, the PRODUCTION OF FLAVONOL GLYCOSIDES (PFG) family of R2R3-MYB transcription factors has a pivotal function in the regulation of genes involved in the phenylpropanoid biosynthetic pathway leading to flavonol synthesis and accumulation (Stracke et al., 2007). In contrast to *CHS* regulation, however, *FAH1* expression was not dependent on PFG1/MYB12, PFG2/MYB11 and PFG3/MYB111, (Supplemental Figure S2B, C) (Stracke et al., 2010). Collectively, our data show that *FAH1* expression is induced under UV-B, and this response is dependent on UVR8 as well as HY5 and HYH.

**Figure 2.**
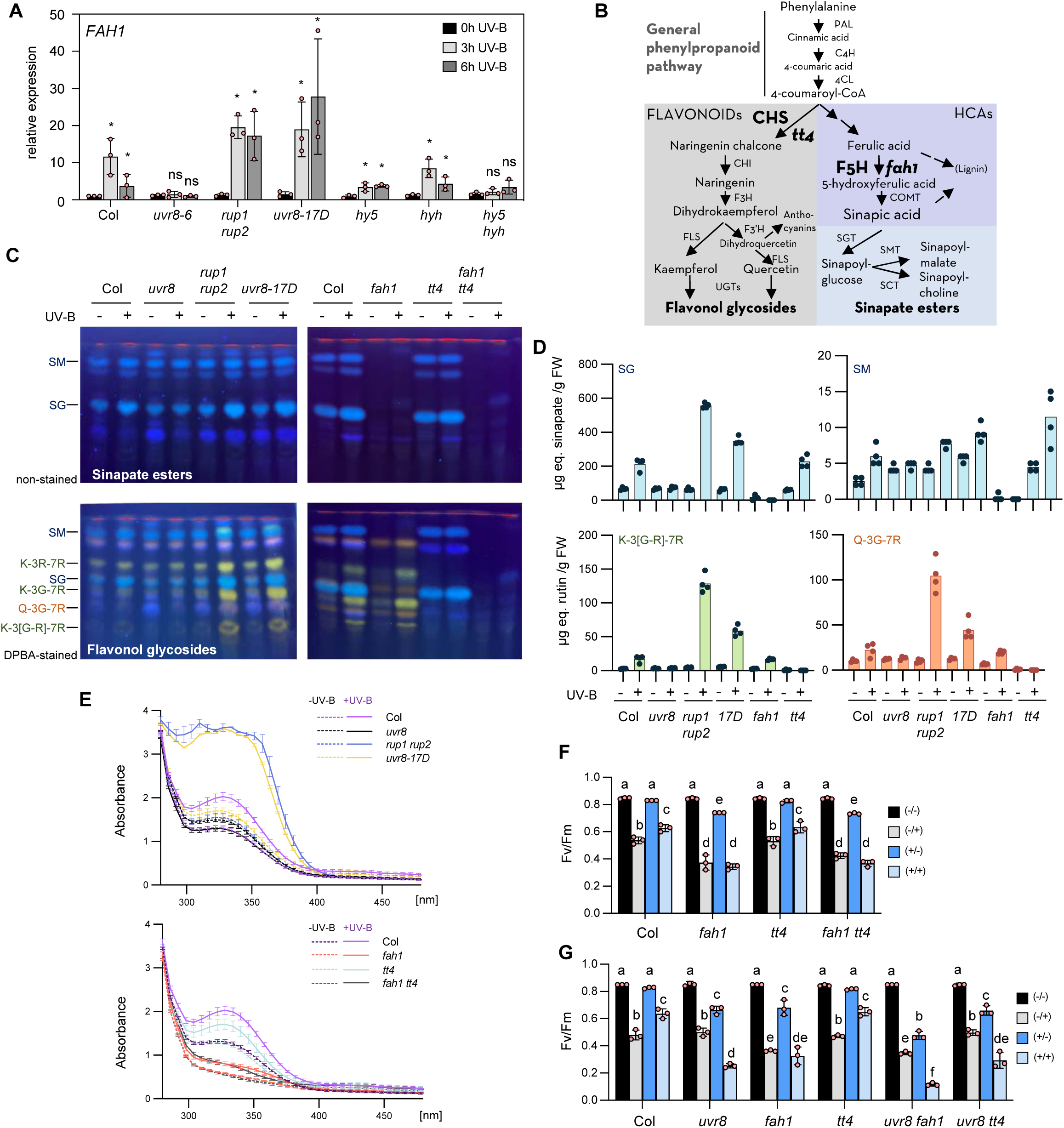
UVR8 signalling induces *FAH1* expression and accumulation of sinapate esters underlying UV-B acclimation. **(A)** qRT-PCR analysis of *FAH1* gene expression changes in response to 3h and 6h supplemental UV-B (0.08 mW cm^-2^) in 7-d-old light grown (20 µmol m^-2^ s^-1^) *uvr8-6* (*uvr8*), *rup1 rup2*, *uvr8-17D*, *hy5*, *hyh*, and *hy5 hyh* seedlings compared to wild type (Col). Values of independent measurements and means ± SD are shown (*n* = 3). * indicate significant difference between the means of UV treated (3 or 6h) and not UV treated (0h) (P< 0.05), as determined by two-way ANOVAs on log2 transformed values. **(B)** Simplified schematic of the phenylpropanoid biosynthetic pathway towards sinapate esters and flavonol glycosides in Arabidopsis. CHS and F5H are highlighted as the key enzymes for synthesis of flavonol glycosides and sinapate esters, respectively, with *tt4* and *fah1* as the respective corresponding mutants. PAL, phenylalanine ammonium lyase ; C4H, cinnamate-4-hydroxylase ; 4CL, 4-coumarate:CoA ligase ; CHS, chalcone synthase ; CHI, chalcone isomerase ; F3H, flavanone 3-hydroxylase ; FLS, flavonol synthase ; F3’H, flavonoid 3′-hydroxylase ; UGTs, UDP-dependent glycosyltransferase ; F5H, FERULIC ACID 5-HYDROXYLASE 1 ; COMT, CAFFEATE O-METHYLTRANSFERASE ; SGT, Sinapate:UDP-glucose glucosyltransferase ; SMT, sinapoylglucose:malate sinapoyltransferase ; SCT, sinapoylglucose:choline sinapoyltransferase **(C)** HPTLC analysis of sinapate ester (*upper panels*) and flavonol glycoside (*lower panels*) levels in 10-d-old seedlings of wild type (Col), *uvr8-6* (*uvr8*), *rup1 rup2*, *uvr8-17D*, *fah1-101* (*fah1*), *tt4*, and *fah1-101 tt4* (*fah1 tt4*) grown for 7 days in white light (20 µmol m^-2^ s^-1^) and exposed to supplemental UV-B (0.08 mW cm^-2^) for 3 days (UV-B: +) or not (UV-B: -). SM, sinapoyl malate; K-3R-7R, kaempferol-3-*O*-rhamnoside-7-*O*-rhamnoside; SG, sinapoyl glucose; K-3G-7R, kaempferol-3-*O*-glucoside-7-*O*-rhamnoside; Q-3G-7R, quercetin-3-*O*-glucoside-7-*O*-rhamnoside; K-3[G-R]-7R, kaempferol 3-*O*-[rhamnosyl-glucoside]-7-*O*-rhamnoside. **(D)** LC-MS-based targeted quantification of the secondary metabolites sinapoyl glucose (SG), sinapoyl malate (SM), kaempferol 3-*O*-[rhamnosyl-glucoside]-7-*O*-rhamnoside (K-3[G-R]-7R), and quercetin-3-*O*-glucoside-7-*O*-rhamnoside (Q-3G-7R) in 10-d-old seedlings of wild type (Col), *uvr8-6* (*uvr8*), *rup1 rup2*, *uvr8-17D* (*17D*), *fah1-101* (*fah1*), and *tt4* grown for 7 days in white light (20 µmol m^-2^ s^-1^) and exposed to supplemental UV-B (0.08 mW cm^-2^) for 3 days (UV-B: +), or not (UV-B: -). Values of independent samples and means are shown (*n* = 4). μg eq. sinapate, μg equivalent of sinapate; μg eq. rutin, μg equivalent of rutin; FW, fresh weight. **(E)** Absorption spectra (280-480 nm) of methanolic extracts extracts (2:1, v/FW) from 10-d-old seedlings of wild type (Col), *uvr8-6* (*uvr8)*, *rup1 rup2*, *uvr8-17D*, *fah1-101* (*fah1*), *tt4* and *fah1-101 tt4* (*fah1 tt4*) grown for 7 days in white light (20 µmol m^-2^ s^-1^) and exposed to supplemental UV-B (0.08 mW cm^-2^) for 3 days (+UV, purple bars) or not (-UV, grey bars). The inset shows the absorption spectra of Col. Graph bar values were calculated as total area under the curves for each genotype and conditions, between 280 and 400 nm, from independent absorption spectra measurements. Means ± SEM are shown (*n* = 3). * indicate significant difference (P < 0.05) between -UV and +UV measurements of same genotype (two-way ANOVAs followed by Bonferroni’s test). FW, fresh weight. **(F, G)** Fv/Fm measurements of 10-d-old seedlings of (E) wild type (Col), *fah1-101 (fah1)*, *tt4* and *fah1-101 tt4* (*fah1 tt4*), and (F) wild type (Col), *uvr8-6* (*uvr8)*, *fah1-101* (*fah1*), *tt4, uvr8-6 fah1-101* (*uvr8 fah1*), and *uvr8-6 tt4* (*uvr8 tt4*) grown as described in Figure 1a. Values of independent experiments and means ± SD are shown (*n* = 3). Shared letters indicate no statistically significant difference between the means (P > 0.05), as determined by two-way ANOVAs followed by Tukey’s test for multiple comparisons. -/-, not acclimated, not stressed; -/+, not acclimated, stressed; +/-, acclimated, not stressed; +/+, acclimated, stressed.

We next analysed UVR8-dependent activation of the phenylpropanoid biosynthetic pathway at the metabolic level (Figure 2B). Methanolic extracts from UV-B treated plants were analysed by high-performance thin layer chromatography (HPTLC) and quantified by ultra-high performance liquid chromatography–tandem mass spectrometry (UHPLC-MS/MS; see Supplemental Tables S1-S3 for quantification of four HCA esters and four flavonoid compounds). For HPTLC, sinapate esters were visualized by their blue autofluorescence under UV-A, whereas flavonol glycosides were visualized under UV-A after staining of the HPTLC plate with diphenylboric acid 2-aminoethylester (DPBA) (Stracke et al., 2007; Stracke et al., 2010). In agreement with the *FAH1* expression data, we observed increased accumulation of sinapate esters (e.g. sinapoyl glucose, SG) in wild type exposed to UV-B (Figure 2C, *upper panel*, Figure 2D, and Supplemental Table S1). Increased sinapate esters were detectable in addition to the well-known accumulation of flavonol glycosides (i.e. kaempferol and quercetin glycosides) under UV-B (Figure 2C, *lower panel*, Figure 2D, and Supplemental Table S1). Increase in these metabolites under UV-B was dependent on UVR8 as well as HY5 and HYH, as indicated by an absence of UV-B-induced accumulation in *uvr8* and *hy5 hyh* (Figure 2D, E, Supplemental Figure S2D, and Supplemental Table S1). *uvr8-17D* and *rup1 rup2* showed elevated accumulation of flavonol glycosides and sinapate esters (Figure 2C, D), whereas this response was reduced in *uvr8-17D hy5* and absent in *uvr8-17D hy5 hyh* (Supplemental Figure S2D).

Next we used a series of mutants, including *fah1* and the *CHS* knock-out mutant *transparent testa 4 (tt4)*, to dissect the contribution of different phenylpropanoid metabolites to UV photoprotection (Figure 2B). As expected, *fah1* mutants did not accumulate sinapate esters (e.g. sinapoyl glucose and sinapoyl malate) but accumulated wild-type levels of flavonol glycosides, whereas *tt4* mutants did not accumulate flavonol glycosides but accumulated wild-type levels of sinapate esters (Figure 2B-D). Absorption spectra of methanolic extracts showed that UV-absorbing compounds accumulated under UV-B in wild type and even more so in *rup1 rup2* and *uvr8-17D*, but not in *uvr8* and *hy5 hyh* mutants (Figure 2E, and Supplemental Figure S2E). Extracts from the *fah1* mutant showed very low UV-absorption capacity, which only slightly increased following UV-B acclimation (Figure 2E). However, extracts from the *tt4* mutant exhibited better UV absorption, which also further increased upon UV-B acclimation (Figure 2E). In agreement, Fv/Fm measurements showed that *fah1* mutants, in contrast to *tt4*, are hypersensitive to UV stress with no improvement following acclimatory UV-B pre-treatment (Figure 2F, G, and Supplemental Figure S2F, G). Moreover, *tt4* displayed Fv/Fm values comparable to those of wild type, and the *fah1 tt4* double mutant was not more sensitive to UV stress than the *fah1* single mutant (Figure 2F). Altogether, our data suggest that sinapate esters play a dominant role as both basal and UV-B-induced “sunscreen”, thereby providing UV photoprotection in Arabidopsis seedlings.

### Enhanced UVR8 signalling rescues photoprotection in *fah1* mutants, associated with the accumulation of HCA derivatives coumaroyl glucose and feruloyl glucose

We observed that *uvr8 fah1*, but not *uvr8 tt4*, was more sensitive to UV stress than both *uvr8* and *fah1* single mutants (Figure 2G), suggesting that FAH1 and UVR8 may also contribute independently to UV photoprotection. Therefore, we tested whether enhanced UVR8 signalling can suppress *fah1* UV-stress hypersensitivity. Indeed, compared to the *fah1* single mutant, enhanced UVR8 activity in *uvr8-17D fah1* and *rup1 rup2 fah1* strongly improved photoprotection specifically under UV-B acclimation conditions (+/+, Figure 3A). Strikingly, the triple mutant *uvr8-17D fah1 tt4* was able to acclimate and induce photoprotection under UV-B comparable to wild type (Figure 3A). Thus, enhanced UVR8 signalling provided by the UVR8^G101S^ variant expressed in *uvr8-17D* can induce a FAH1- and CHS-independent photoprotection mechanism. However, the absorbance spectra of *uvr8-17D fah1* and *uvr8-17D fah1 tt4* methanolic extracts indicated that elevated UV-absorbing metabolites were present despite the absence of FAH1 and CHS (Figure 3B, C). This suggests that enhanced UVR8 signalling in the *uvr8-17D* background is associated with UV-B acclimation even in the absence of sinapoyl malate, sinapoyl glucose, and flavonol glycosides (Figure 3C, and Supplemental Table S2). Intriguingly, HPTLC analysis of methanolic extracts from UV-B-acclimated *uvr8-17D fah1* and *uvr8-17D fah1 tt4* revealed an additional, prominent band under non-stained conditions (highlighted with an asterisk in Figure 3C). This band was also detectable, although weakly, in UV-B-acclimated extracts of *fah1*, *fah1 tt4*, and *uvr8-17D tt4* (Figure 2C and Figure 3C). We isolated the band in question directly from a HPTLC plate carrying extracts of UV-B-treated *uvr8-17D fah1 tt4*, as well as a wild-type control from the corresponding position on the HPTLC plate. UHPLC-MS/MS was then used and identified its constituents as the HCA derivatives coumaroyl glucose and feruloyl glucose (Supplemental Figure S3). These two metabolites were indeed able to absorb UV (Supplemental Figure S3A, B). Targeted LC-MS analysis confirmed strong accumulation of coumaroyl glucose in UV-B-acclimated seedlings expressing hyperactive UVR8^G101S^ (*uvr8-17D*, *uvr8-17D fah1*, *uvr8-17D tt4* and, particularly, *uvr8-17D fah1 tt4*) (Figure 3D, see also Supplemental Table S2 for quantification of other HCA esters and flavonol glycosides), whereas feruloyl glucose seemed to accumulate only in UV-B-acclimated seedlings wherein FAH1 was absent (*fah1*, *uvr8-17D fah1*, and, particularly, *uvr8-17D fah1 tt4*) (Figure 3E, and Supplemental Table S2). Thus, UV-B acclimation and UV tolerance in *uvr8-17D fah1 tt4* is specifically associated with the accumulation of UV-absorbing coumaroyl glucose and feruloyl glucose. In addition, this may contribute to the hypersensitivity of *uvr8 fah1* compared to *fah1* (Figure 2G), as the induction of the feruloyl and coumaroyl glucose is UV-B-mediated and therefore absent in a *uvr8* mutant (Supplemental Table S2).

**Figure 3.**
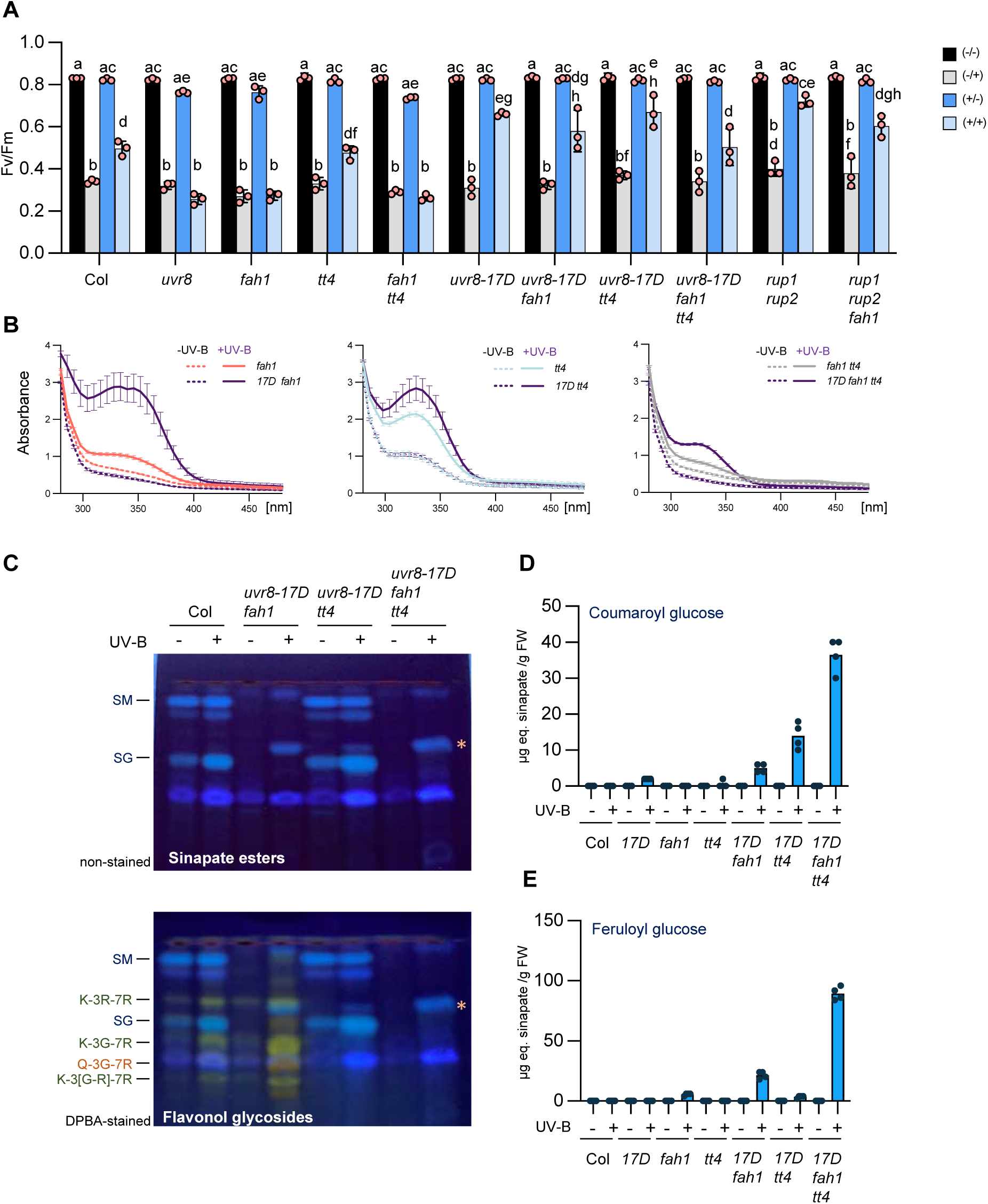
Enhanced UVR8 signalling rescues *fah1* photoprotection and promotes accumulation of unique HCAs. **(A)** Fv/Fm measurements of 10-d-old seedlings of wild type (Col), *uvr8-6* (*uvr8*), *fah1-101*(*fah1*), *tt4*, *fah1-101 tt4* (*fah1 tt4*), *uvr8-17D*, *uvr8-17D fah1-101* (*uvr8-17D fah1*), *uvr8-17D tt4, uvr8-17D fah1-101 tt4* (*uvr8-17d fah1 tt4*), *rup1 rup2* and *rup1 rup2 fah1-101* (*rup1 rup2 fah1*) grown as described in Figure 1a. Values of independent experiments and means ± SD are shown (*n* = 3). Shared letters indicate no statistically significant difference between the means (P > 0.05), as determined by two-way ANOVAs followed by Tukey’s test for multiple comparisons. -/-, not acclimated, not stressed; -/+, not acclimated, stressed; +/-, acclimated, not stressed; +/+, acclimated, stressed. **(B)** Absorption spectra (280-480 nm) of methanolic extracts extracts (2:1, v/FW) from 10-d-old seedlings of *uvr8-17D* (*17D*), *uvr8-17D fah1-101* (*17D fah1*), *tt4*, *uvr8-17D tt4* (*17D tt4*), and *uvr8-17D fah1-101 tt4* (*17d fah1 tt4*) grown for 7 days in white light and exposed to supplemental UV-B for 3 days (+UV-B, purple lines) or not (-UV-B, grey lines). Data of *fah1 tt4* are identical to the ones in figure 2D (all samples were analysed in parallel). Values of independent measurements and means ± SEM are shown (*n* = 3). **(C)** HPTLC analysis of sinapate esters (*upper panels*) and flavonol glycosides (*lower panels*) levels in 10-d-old seedlings of wild type (Col), *uvr8-17D fah1-101*, *uvr8-17D tt4*, and *uvr8-17D fah1-101 tt4* grown for 7 days in white light (20 µmol m^-2^ s^-1^) and exposed to supplemental UV-B (0.08 mW cm^-2^) for 3 days (UV-B: +) or not (UV-B: -). SM, sinapoyl malate; K-3R-7R, kaempferol-3-*O*-rhamnoside-7-*O*-rhamnoside; SG, sinapoyl glucose; K-3G-7R, kaempferol-3-*O*-glucoside-7-*O*-rhamnoside; Q-3G-7R, quercetin-3-*O*-glucoside-7-*O*-rhamnoside; K-3[G-R]-7R, kaempferol 3-*O*-[rhamnosyl-glucoside]-7-*O*-rhamnoside. The asterisk (*) indicates coumaroyl glucose and feruloyl glucose, as identified by LC-MS/MS (see Supplemental Figure S4). **(D, E)** Quantification of (E) coumaroyl glucose and (F) feruloyl glucose by LC-MS-based analysis of methanolic extracts from 10-d-old seedlings of wild type (Col), *uvr8-17D* (*17D*), *fah1-101* (*fah1*), *tt4*, *uvr8-17D fah1-101*, *uvr8-17D tt4*, and *uvr8-17D fah1-101 tt4* grown for 7 days in white light (20 µmol m^-2^ s^-1^) and exposed to supplemental UV-B (0.08 mW cm^-2^) for 3 days (UV-B: +) or not (UV-B: -). Values of independent samples and means are shown (*n* = 4). μg eq. sinapate, μg equivalent of sinapate; FW, fresh weight.

### FAH1 plays a major role for plant performance in presence of UV-B

Our data indicate that FAH1 and sinapate ester accumulation are key for photoprotection against UV stress at the seedling stage. We thus further examined plant growth and survival for *fah1* and *tt4* mutants under UV. On soil, in the absence of UV-B, *uvr8*, *fah1*, *tt4*, and *fah1 tt4* grew and developed similarly to wild type (Figure 4A). However, under supplemental UV-B, *uvr8*, *fah1*, and *fah1 tt4* were affected and showed strongly impeded growth and damaged PSII in comparison to wild type and *tt4*, which were less impaired (Figure 4A, B), supporting that induced FAH1 and sinapate ester accumulation plays a major role for photoprotection against UV-B. TT4 seemed to play a less important but clear role in development under UV-B. Indeed, the *fah1 tt4* rosettes were smaller than *fah1* rosettes after 4 weeks under supplemental UV-B and the Fv/Fm ratio was lower (Figure 4A, B). In agreement with the Fv/Fm data of seedlings, UVR8^G101S^–mediated hyperactivation of UV-B signalling in *uvr8-17D fah1* and *uvr8-17D fah1 tt4* suppressed the UV stress hypersensitivity of *fah1* and *fah1 tt4*, resulting in less photoinhibition (Figure 4B). It is of note, however, that reduced growth of *uvr8-17D* lines under UV-B is due to enhanced UV-B-induced photomorphogenesis (Podolec et al., 2021b), and is not representing a UV-B stress phenotype – in contrast to *uvr8*, *fah1*, and *fah1 tt4* (Figure 4B).

**Figure 4.**
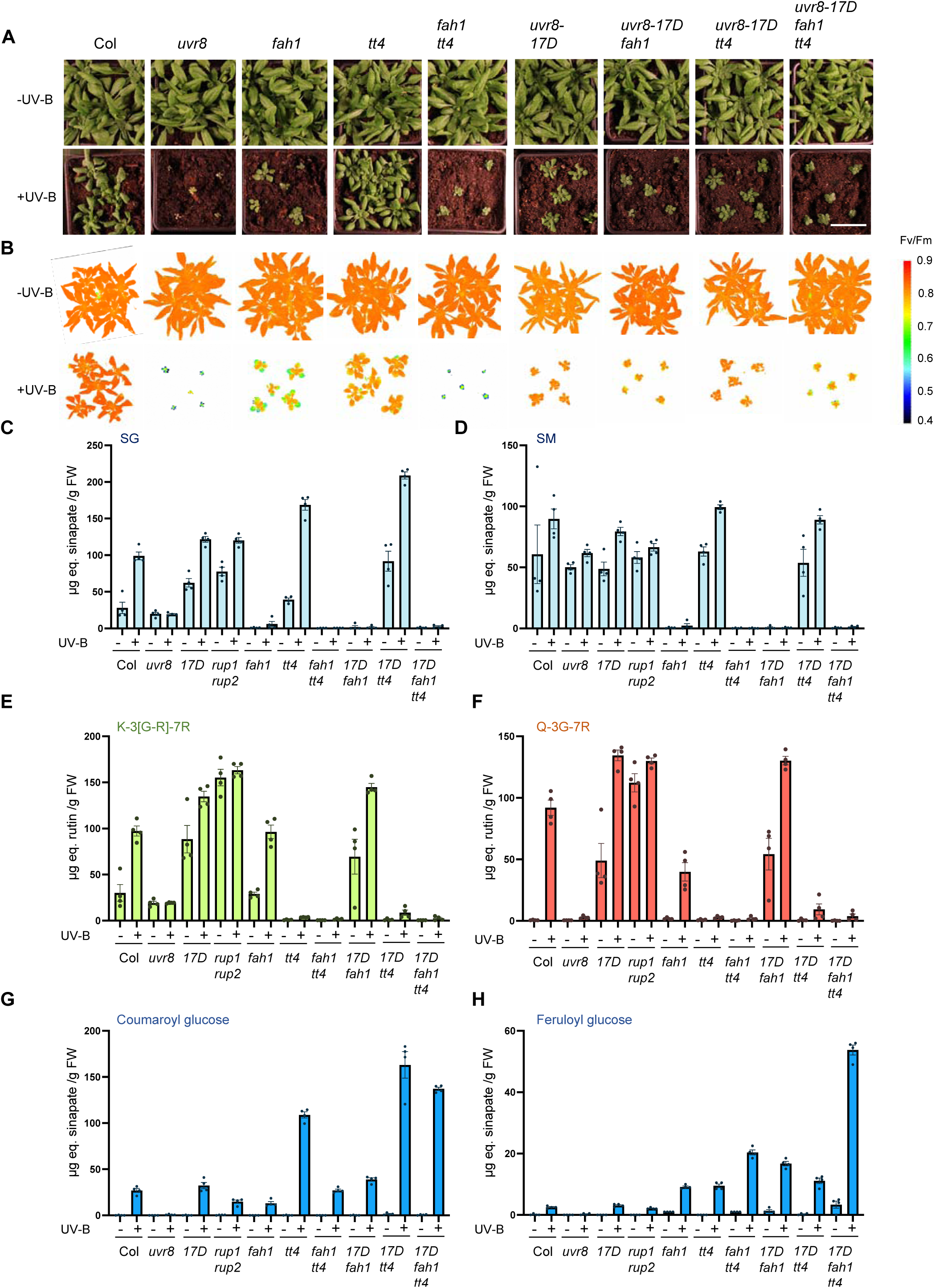
FAH1 plays a major role for photoprotection in soil-grown plants. **(A)** Photographs of 4-week-old wild-type (Col), *uvr8-6* (*uvr8*), *fah1-101* (*fah1*), *tt4*, *fah1-101 tt4* (*fah1 tt4*), *uvr8-17D*, *uvr8-17D fah1-101* (*uvr8-17D fah1*), *uvr8-17D tt4*, and *uvr8-17D fah1-101 tt4* (*uvr8-17D fah1 tt4*) plants grown under long-day conditions under 100 µmol m^-2^ s^-1^ of white light with supplemental UV-B (+UV-B, 0.3 mW cm^-2^, *lower panels*) or not (-UV-B, *upper panels*). Bar = 4cm. **(B)** False-color image representing Fv/Fm values of wild-type (Col), *uvr8-6* (*uvr8*), *fah1-101*(*fah1*), *tt4*, *fah1-101 tt4* (*fah1 tt4*), *uvr8-17D*, *uvr8-17D fah1-101* (*uvr8-17D fah1*), *uvr8-17D tt4*, and *uvr8-17D fah1-101 tt4* (*uvr8-17D fah1 tt4*) plants grown under long-day conditions under 100 µmol m^-2^ s^-1^ of white light with supplemental UV-B (+UV-B, 0.3 mW cm^-2^, *lower panels*) or not (-UV-B, *upper panels*). **(C-H)** Accumulation of selected HCA esters and flavonol glycosides in soil grown plants. LC-MS-based targeted metabolite quantifications of (C) sinapoyl glucose (SG), D) sinapoyl malate (SM), (E) kaempferol 3-*O*-[rhamnosyl-glucoside]-7-*O*-rhamnoside (K-3[G-R]-7R), (F) quercetin-3-*O*-glucoside-7-*O*-rhamnoside (Q-3G-7R), (G) coumaroyl glucose, and (H) feruloyl glucose in 3-week-old wild type (Col), *uvr8-6* (*uvr8*), *uvr8-17D* (*17D*), *rup1 rup2*, *fah1-101* (*fah1*), *tt4*, *fah1-101 tt4* (*fah1 tt4*), *uvr8-17D fah1-101* (*17D fah1*), *uvr8-17D tt4* (*17D tt4*) and *uvr8-17D fah1-101 tt4* (*17D fah1 tt4*) grown on soil under long-day conditions under 100 µmol m^-2^ s^-1^ of white light in the absence of UV-B and exposed (+) or not (-) to supplemental UV-B (0.3 mW cm^-2^) for 24 hours. Values of independent samples and means ± SEM are shown (*n* = 4). μg eq. sinapate, μg equivalent of sinapate; μg eq. rutin, μg equivalent of rutin; FW, fresh weight.

Targeted metabolite profiling of methanolic extracts from 3-week-old soil-grown plants confirmed UVR8-dependent accumulation of sinapate esters, particularly sinapoyl glucose in wild type and *tt4* plants, and its absence in *fah1* mutants (Figure 4C, D, and Supplemental Table S3 for quantification of other HCA esters and flavonol glycosides). Conversely, UVR8-dependent accumulation of flavonol glycosides was detectable in *fah1* but not in *tt4* mutants (Figure 4E, F). Interestingly, as seen in *in vitro*-grown seedlings (Figure 3), the increased UV tolerance of *uvr8-17D fah1* and *uvr8-17D fah1 tt4* versus that of *fah1* and *fah1 tt4* was associated with the over-accumulation of coumaroyl glucose and feruloyl glucose compared to wild-type (Figure 4G, H). We conclude that UVR8-induced accumulation of sinapate esters, or derivatives of its precursors in absence of F5H activity in *fah1* mutants, plays a major role in UV-B acclimation in soil-grown plants, and together with flavonols ensure plant survival under UV.

### Photoreceptor signalling pathways converge on *FAH1* expression and act in concert for UV photoprotection

Phytochrome and cryptochrome signalling contribute to UV tolerance (Rai et al., 2019; Tissot and Ulm, 2020; Stockenhuber et al., 2024). In agreement, under prolonged UV exposure, *uvr8* mutants are more UV sensitive than wild type, but *uvr8 cry1* and *uvr8 phyB* are even more UV sensitive than *uvr8* mutants (Figure 5A) (Tissot and Ulm, 2020). By contrast, *cry1 phyB* are less affected in presence of UV-B (Figure 5A) (Tissot and Ulm, 2020). Strikingly, the *uvr8 cry1 phyB* triple mutant was even more UV sensitive than *uvr8 cry1* and *uvr8 phyB* double mutants (Figure 5A). Indeed, *uvr8 cry1 phyB* survived under white light but died after about 6 weeks under white light supplemented with UV under conditions where even *uvr8 cry1* and *uvr8 phyB* survived (Figure 5A). In agreement, analysis of non-acclimated plants under UV stress confirmed that cry1 and phyB plays a role in basal UV protection, as Fv/Fm values for non-acclimated *cry1*, *uvr8 phyB*, *uvr8 cry1*, *cry1 phyB,* and *uvr8 cry1 phyB* under UV stress were lower than those for wild type (Figure 5B). In agreement, visible light induced *FAH1* expression was impaired in *cry1* and *phyB* mutant backgrounds (Figure 5C). These findings indicate that photoreceptors for blue light, red light, and UV-B cooperatively regulate UV photoprotection, activating *FAH1* expression, and accumulating key UV-absorbing compounds.

**Figure 5.**
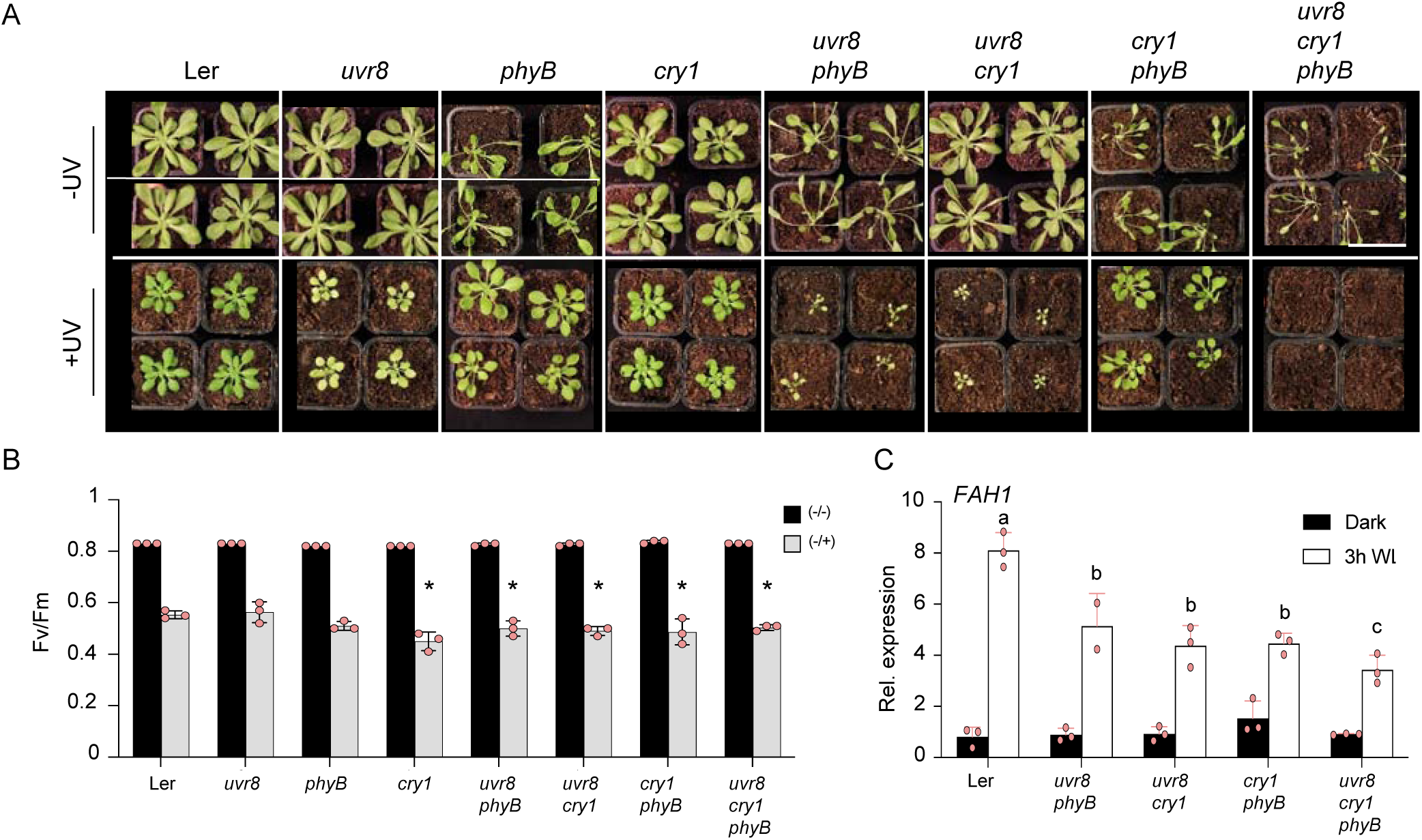
Cryptochromes and phytochromes redundantly contribute to UVR8-mediated *FAH1* expression and UV-B tolerance. **(A)** Photographs of 6-week-old wild-type (Ler), *uvr8-1*, *phyB-5*, *hy4-2.23N* (*cry1*), *uvr8-1 phyB-5* (*uvr8 phyB*), *uvr8-1 hy4-2.23N* (*uvr8 cry1*), *hy4-2.23N phyB-5* (*cry1 phyB*), and *uvr8-1 hy4-2.23N phyB-5* (*uvr8 cry1 phyB*) plants grown under short-day conditions with 100 µmol m^-2^ s^-1^ of white light with supplemental UV-B (+UV, 0.06 mW cm^-2^) or not (-UV). Bars = 5cm **(B)** Fv/Fm measurements of 7-d-old seedlings of wild-type (Ler), *uvr8-1*, *phyB-5*, *hy4-2.23N* (*cry1*), *uvr8-1 phyB-5* (*uvr8 phyB*), *uvr8-1 hy4-2.23N* (*uvr8 cry1*), *hy4-2.23N phyB-5* (*cry1 phyB*), and *uvr8-1 hy4-2.23N phyB-5* (*uvr8 cry1 phyB*) plants grown under long-day conditions with 100 µmol m^-2^ s^-1^ of white light without supplemental UV-B and exposed (-/+) or not (-/-) to broad-band UV-B for 2 hours starting at ZT4. Values of independent experiments and means ± SD are shown (*n* = 3). Asterisks indicate statistically significant difference between the means (p < 0.05), as determined by two-way ANOVAs followed by Tukey’s test for multiple comparisons. -/-, not acclimated, not stressed; -/+, not acclimated, stressed. **(C)** qRT-PCR of *FAH1* expression in 7-d-old seedlings of wild-type (Ler), *uvr8-1* (*uvr8*), *uvr8-1 phyB-5* (*uvr8 phyB*), *uvr8-1 hy4-2.23N* (*uvr8 cry1*), *hy4-2.23N phyB-5* (*cry1 phyB*), and *uvr8-1 hy4-2.23N phyB-5* (*uvr8 cry1 phyB*) grown for 4 days under long-day conditions with 100 µmol m^-2^ s^-1^ of white light without supplemental UV-B and dark adapted for 3 days, then exposed for 3h to white light (100 µmol m^-2^ s^-1^) or kept in darkness (Dark). Values of independent measurements and means ± SD are shown (*n* = 3). Shared letters indicate no statistically significant difference between the means (P > 0.05), as determined by two-way ANOVAs followed by Tukey’s test for multiple comparisons.

## DISCUSSION

UVR8 and cryptochromes act together to ensure plant survival when grown in the field in the presence of UV (Rai et al., 2019; Stockenhuber et al., 2024). However, the extent to which other photoprotective mechanisms activated upon light perception contribute to UV tolerance is not well understood. Our findings suggest that photoreceptor-mediated regulation of biosynthesis of UV-absorbing sinapate esters is a key response to ensure plant survival under UV.

Sinapate esters belong to the family of phenylpropanoids which comprises specialized metabolites with broad functions including roles in pollinator attraction, cuticle composition, pathogen defence, and photoprotection against UV and high light (Sheahan, 1996; Rozema et al., 2009; Erb and Kliebenstein, 2020; Ferreyra et al., 2021). The phenylpropanoid pathway diverges at 4-coumaroyl-CoA, providing flavonoids (incl. flavonols and anthocyanins) and HCAs like sinapate, as well as its conjugated forms (Fraser and Chapple, 2011). Flavonoids are generally known to be photoreceptor-inducible and are thus the focus of many studies in the field, whereas sinapate esters are usually considered constitutively present and often neglected in the literature (Lois, 1994; Kliebenstein et al., 2002; Morales et al., 2010; Stracke et al., 2010; Kusano et al., 2011; Neugart et al., 2019; Rai et al., 2019; Podolec et al., 2021a), despite the fact that they are highly efficient to absorb across a broad range of UV wavelength (Dean et al., 2014). However, UVR8-induced sinapate ester accumulation was described in response to UV-A (Brelsford et al., 2019) and in the context of UV-B-induced defense responses against the necrotrophic fungal pathogen *Botrytis cinerea* (Demkura and Ballare, 2012). Moreover, transgenic enhancement of sinapate synthesis was also found to enhance the resistance to diamondback moth (*Plutella xylostella*) herbivory (McInnes et al., 2023). Our data show that, in addition to the induction of flavonoid biosynthesis, UVR8, cryptochromes and phytochromes indeed activate *FAH1* expression through the HY5 and HYH transcription factors, promoting sinapate ester accumulation and photoprotection. HY5 functions often in concert with BBX factors, such as BBX20–22, BBX29, and BBX31, that were shown to promote accumulation of soluble phenylpropanoid derivatives and may thus contribute to UV-B acclimation (Yadav et al., 2019; Bursch et al., 2020; Xu, 2020; Podolec et al., 2022; Medina-Fraga et al., 2023). Whether *FAH1* expression can be modulated by other transcriptional regulators associated with UVR8 signalling remains to be determined (Liang et al., 2018; Liang et al., 2019; Liang et al., 2020; Qian et al., 2020; Tavridou et al., 2020a; Tavridou et al., 2020b; Yang et al., 2020; Podolec et al., 2021a). Moreover, other transcription factors presently not linked to UVR8 signalling may additionally modulate the response leading to sinapate esters at different levels. For example, MYB4 represses expression of *C4H*, encoding cinnamate 4-hydroxylase, and *myb4* mutants show enhanced levels of sinapate esters and enhanced tolerance to UV-B stress (Jin et al., 2000). Downregulation of *MYB4* in response to UV-B may thus contribute to sinapate ester accumulation and enhanced UV-absorbing capability (Jin et al., 2000).

We used chlorophyll fluorescence measurement as a non-invasive approach and calculated the Fv/Fm to assess PSII integrity, thereby providing an estimation of PSII damage. As we measured the Fv/Fm directly after UV stress treatment, the observed decrease likely reflects the direct damage caused by UV on D1 and the oxygen evolving center (Takahashi et al., 2010). Absorption spectra and physiological analyses of WT and *fah1* seedlings and rosette-stage plants show that sinapate esters provide UV-screening properties in plant extracts, make a major contribution to direct photoprotection of the photosynthetic machinery upon UV-stress treatment, and play a major role for plant survival under UV in lab conditions. Flavonoids seemed to play a lesser role in UV-B-induced acclimation and UV tolerance under our experimental conditions, importantly at both *in vitro* seedling and more mature soil grown plant stages as indicated by less pronounced UV-induced damage at PSII in *tt4* compared to *fah1* and much better growth under UV; however, the small rosette phenotype of *tt4* and additive effect of *tt4* and *fah1* mutations after 3 weeks under supplemental UV-B indicates that flavonoids contribute to plant performance under UV, in agreement with previous reports (Landry et al., 1995; Stracke et al., 2010). Thus, although flavonoids are diverse, abundant, and highly UV-inducible, the function of HCA-derived compounds seems of even greater importance for UV tolerance, at least under the laboratory conditions used in this work. As the plant extracts derived from *tt4* mutants absorb UV relatively well, it is possible that, more than UV-absorbing activity, the antioxidant or visible light–absorbing properties of flavonoids may contribute to UV tolerance, complementary to sinapate esters UV-screen function (Agati and Tattini, 2010; Hideg et al., 2013). However, it has also previously been reported that absence of CHS, and thus flavonoids, has no detectable effect on plant phenotype under prolonged UV exposure in a sun simulator, further supporting that HCAs might indeed play a more important role than flavonoid to ensure plant survival under more natural conditions in presence of UV (Rai et al., 2019). As we observed that combinatorial *uvr8*, *cry1*, and *phyB* mutants are affected for *FAH1* expression, and cannot survive under prolonged UV exposure, we propose that induction of *FAH1* and associated sinapate synthesis is a key mechanism for UV tolerance, although we do not exclude that other photoreceptor-induced photoprotective mechanisms contribute to UV tolerance, which may even be of much higher relative importance under other growth conditions. Future experiments in the field will be of great interest and importance to further characterize the contribution of flavonoids and sinapate esters in a more complex light environment, and the influence of other environmental factors on light-and UV-induced plant photoprotection.

The development of UV-B protective mechanisms has played a pivotal role in the successful terrestrialization of plants and their colonization of land (Rozema et al., 1997). UV inducibility of sunscreen capacity is broadly spread across the plant kingdom, although the UV-absorbing compounds may differ. The UVR8 photoreceptor is well-conserved across the green lineage (Rizzini et al., 2011; Jenkins, 2014; Han et al., 2019; Podolec et al., 2021a; Zhang et al., 2022), with functional homologs of Arabidopsis UVR8 characterised in the single-cell green algae *Chlamydomonas reinhardtii* (Allorent et al., 2016; Tilbrook et al., 2016); in bryophytes, namely in the liverwort *Marchantia polymorpha* and the moss *Physcomitrium patens* (Soriano et al., 2018; Kondou et al., 2019); and in the flowering plant tomato (*Solanum lycopersicum*) (Li et al., 2018; Liu et al., 2020). UV-B-mediated activation of phenylpropanoid biosynthesis and corresponding gene expression appears to be broadly conserved among land plants (Stracke et al., 2010; Wolf et al., 2010; Schreiner et al., 2017; Clayton et al., 2018; Ferreyra et al., 2021). Phenylpropanoid compounds are not present in bacteria, but the accumulation of UV-absorbing, mycosporin-like amino acids was shown to be UV-B-inducible in cyanobacteria (Rozema et al., 2002; Sinha et al., 2002). Sinapate esters accumulate to a high level in Brassicaceae and have also been detected in other plant species (Milkowski and Strack, 2010; Nguyen et al., 2021). A natural variation study reported Arabidopsis accessions that accumulate high levels of phenylacylated-flavonols (saiginols, including a flavonol-glycoside esterified with sinapoyl) in floral tissues, a trait which was associated with greater UV tolerance and was primarily found in accessions prevalent in regions with high UV-B irradiance (Tohge et al., 2016; Tohge and Fernie, 2017). Evolution may have selected for a diversification of phenylpropanoid-derived compounds towards better adaptation to the light environment (Tohge et al., 2013; de Vries et al., 2021). Such revelations also suggest that genetic variation can be used to increase plant UV tolerance. By combining mutations that enhance UVR8 signalling with those that block synthesis of flavonoids and sinapates, we generated plants that accumulated high levels of UV-absorbing HCAs, namely coumaroyl glucose and feruloyl glucose. An improved understanding of the breadth and relative importance of UV-absorbing metabolites as “sunscreens”, as well as regulation of their synthesis, offers potential opportunities for plant engineering towards better UV tolerance.

## MATERIAL AND METHODS

### Plant material

The Arabidopsis accessions Columbia (Col-0, herein Col) or Landsberg *erecta* (L*er*-0, herein Ler) were used as wild-type controls in all experiments, as indicated. The following published mutants were used in this study: *uvr8-6* (Favory et al., 2009), *uvr8-17D* (Podolec et al., 2021b), *rup1-1 rup2-1* (Gruber et al., 2010), *hy5-215* (Oyama et al., 1997), *hyh*, *hy5-215 hyh* (Zoulias et al., 2020), *fah1-2* (Chapple et al., 1992), *fah1-101* (Maruta et al., 2014), *tt4-11* (Bowerman et al., 2012), *phyA-211* (Nagatani et al., 1993), *phyB-9*, *phyA-211 phyB-9* (Enderle et al., 2017), *myb12*, *myb111*, *myb12 myb111*, and *myb11 myb12 myb111* (Stracke et al., 2007) are all in the Col background, whereas *uvr8-1* (Kliebenstein et al., 2002), *fah1-7* (Meyer et al., 1996), *phyB-5* (Reed et al., 1994), *hy4-2.23N* (Koornneef et al., 1980), and *phyB-5 hy4-2.23N* (Tissot and Ulm, 2020) are in the Ler background. The *fah1-101 tt4-11*, *uvr8-6 fah1-101*, *uvr8-6 tt4-11*, *uvr8-17D fah1-101*, and *uvr8-17D tt4-11* double mutants were generated by crossing the respective single mutants; *uvr8-17D fah1-101 tt4-11* was generated by crossing *uvr8-17D fah1-101* and *uvr8-17D tt4-11*; *uvr8-17D hy5-215*, *uvr8-17D hyh*, and *uvr8-17D hy5-215 hyh* were generated by crossing *uvr8-17D* and *hy5-215 hyh*; and *uvr8-1 hy4-2.23N phyB-5* was generated by crossing *uvr8-1 hy4-2.23N* and *uvr8-1 phyB-5*. Genotyping of *hy5-215*, *rup1-1*, *rup2-1*, *uvr8-6* (Gruber et al., 2010), *uvr8-17D* (Podolec et al., 2021b), and *phyB-5* (Neff and Chory, 1998) was performed as previously published. Supplemental Table S4 lists the primers used for genotyping all other mutants used to generate combinatorial mutants.

### Growth conditions and light treatments

For experiments under aseptic conditions *in vitro*, Arabidopsis seeds were surface sterilized with chlorine gas and sown on half-strength Murashige & Skoog (MS) basal salt medium (Duchefa) containing 0.8% (w/v) agar (Applichem) supplemented with 1% (w/v) sucrose. After 2-day stratification at 4°C, seedlings were grown in white light, or seeds were exposed to 6 h white light (60 µmol m^−2^ s^−1^) before transferring them into darkness at 22°C for experiments involving etiolated seedlings. Except otherwise indicated, seedlings were grown in continuous white light (white-light fluorescent tubes Osram L18W/30, 20 µmol m^−2^ s^−1^), supplemented with narrowband UV-B tubes (Philips TL20W/01RS, 0.08 mW cm^-2^). For UV-stress treatment, plates were irradiated under broadband UV-B tubes (Philips TL20W/12RS; 2.2 mW cm^-2^). For UV stress treatment on photoreceptor mutants (Figure 5B), seedlings were grown under aseptic conditions in long day conditions with a 16-h day/8-h night cycle (22°C/18°C) in a Percival growth chamber (CLF Climatics) with white-light fluorescent tubes Philips Alto II F17T8/TL841 (100 µmol m^−2^ s^−1^) for 7 days, and UV stress treatment was performed at ZT4, for 2 hours. Each experiment was done using 3 replicate plates per condition, each containing all genotypes tested in parallel. Control (-UV-B, under lid and a Schott WG368 long-pass filter, for 10 days) and treatment (+UV-B, under lid and a Schott WG368 long-pass filter for 7 days then under lid and a Schott WG305 long-pass filter for 3 days) plates were positioned side-by-side in the same light field. Given white light and UV-B irradiances were measured under the lid and filter (WG305 or WG368), as will be reaching the respective plants. For qRT-PCR-based *FAH1* expression analysis of photoreceptor mutants under aseptic conditions, seedlings were grown under long day conditions with a 16-h day/8-h night cycle (22°C/18°C) in a Percival growth chamber (CLF Climatics) with white-light fluorescent tubes Philips Alto II F17T8/TL841 (100 µmol m^−2^ s^−1^) for 4 days, dark adapted for 3 days, and kept in darkness or exposed to white light for 3 h.

For UV tolerance assays on soil, plants were either grown under long-day conditions with a 16-h day/8-h night cycle (22°C/18°C) in a Percival growth chamber (CLF Climatics) with white-light fluorescent tubes Philips Alto II F17T8/TL841 (100 µmol m^−2^ s^−1^) supplemented, or not, with Philips TL40/W01RS narrowband UV-B tubes (0.3 mW cm^-2^) (for phenylpropanoid mutants analysis), or under short-day conditions with a 8-h day/16-h night cycle (22°C/18°C; 100 µmol m^−2^ s^−1^)) supplemented, or not, with Philips TL40/W01RS narrowband UV-B tubes (0.06 mW cm^-2^) (for the analysis of photoreceptor mutants).

Fluence rates of visible light were measured with a LI-250 Light Meter (LI-COR Biosciences). UV-B irradiances were measured with a VLX-3W UV Light Meter equipped with a CX-312 sensor (Vilber Lourmat). See also Supplemental Figure S4 for spectra of the different light sources used, as measured with an Ocean Optics QE65000 spectrometer.

### Immunoblot analysis

Total protein extracts from seedlings were prepared as previously described (Samol et al., 2012). In brief, samples were harvested, frozen in liquid nitrogen, ground, and proteins were extracted by incubating in lysis buffer (100 mM Tris-HCl pH 7.7, 2% (w/v) SDS, 50 mM NaF, and Protease Inhibitor Cocktail (Sigma-Aldrich)) for 30 min at 37°C. After 15 min centrifugation at 4°C at 15’000 *g* the clear supernatants were collected, and total protein concentration was measured with Bicinchoninic Acid solution (BCA, Sigma-Aldrich) following manufacturer’s instructions. Ten-μg protein of each sample were separated by electrophoresis in 15% SDS-polyacrylamide gels and transferred to polyvinylidene difluoride (PVDF) membranes (Roth) for 7 min at 20V using the iBlot 2 Dry Blotting System (ThermoFisher Scientific), before blocking in TBS-T with 5% (w/v) nonfat dry milk for 1 h. Polyclonal anti-D1 (AS05084A, C-terminal, Agrisera) and anti-RbcL (AS03037, Agrisera) were used as primary antibodies. Horseradish peroxidase (HRP)-conjugated anti-rabbit immunoglobulins (Dako A/S) were used as secondary antibodies. Signal was revealed using the ECL Select Western Blotting Detection Reagent (GE Healthcare) and analysed using an Amersham Imager 680 camera system (GE Healthcare).

### Quantitative real-time PCR (qRT-PCR)

Plant total RNA was isolated with the Plant RNeasy kit (Qiagen) including DNase treatment according to the manufacture’s standard protocol. Synthesis of cDNA was performed using the Taqman Reverse Transcription Reagents kit (ThermoFisher Scientific). Each qRT-PCR reaction contained cDNA (equivalent of 3 ng RNA) synthetized with a 1:1 mixture of oligo(dT) primers and random hexamers. qRT-PCR was performed using a PowerUp SYBR Green Master Mix (Thermo Fisher Scientific) on a QuantStudio^TM^ 5 Real-Time PCR System (ThermoFisher Scientific). The gene-specific primers used were 5ʹ-ATG ATG GGG ATG TTG TCG AT-3ʹ and 5ʹ-CGT CCA TGA TGA TTG CTT TG-3ʹ for *FAH1* (AT4G36220); and 5’-AGC TGA TGG ACC TGC AGG CAT CTT GGC-3’ and 5’-TGC ATG TGA CGT TTC CGA ATT GTC GAC-3’ for *CHS* (AT5G13930). The ΔΔCT method (Livak and Schmittgen, 2001) was used to calculate expression values, with *PP2AA3 (PROTEIN PHOSPHATASE 2A SUBUNIT A3*, AT1G13320) as a reference gene (Czechowski et al., 2005). All expression values were normalized against an untreated wild type, which was set to 1. Each experiment was performed by combining three independent biological replicates.

### F_v_/F_m_ measurements

A PSI (Photon Systems Instruments) FluorCam 800 MF was used to assay the maximum quantum efficiency of PSII (F_v_/F_m_) by measuring plant chlorophyll fluorescence. Before measurements, plates (10 to 20 seedlings per genotype per plate) were incubated in darkness for at least 5 min, or plants grown on soil (5 plants per pots) were incubated in darkness for 20 min, which was confirmed to be sufficient to reach steady state under the plant growth conditions used (as recommended by Murchie and Lawson, 2013). Maximum fluorescence in the dark (Fm) was measured using saturating pulses of 2000 µmol m^-2^ s^-1^ (white light, 400-720 nm) for 960 ms, followed by orange-red light (620 nm) detection pulses (10 µs). F_v_/F_m_ was calculated as F_v_/F_m_ = (F_m_ – F_o_) / F_m_, where F_m_ is the maximal fluorescence and F_o_ is the minimal fluorescence in the dark-adapted state (Baker, 2008).

### Extraction of phenolic compounds, HPTLC, and measurements of absorption spectra

HPTLC was used to analyse the phenolic compounds profile, as described previously (Stracke et al., 2007; Stracke et al., 2010). Phenolic compounds were extracted from 50 mg of fresh plant material homogenized using a Silamat S6 (Ivodar Vivadent) in 100 μL of 80% (v/v) methanol, incubated for 10 min at 70°C at 800 *g*, and centrifuged for 10 min at 12’000 *g* at room temperature (RT). Forty-µL of supernatant were spotted using capillary tubes on silica-60 HPTLC glass plates (Sigma-Aldrich) used as the stationary phase. Adsorption chromatography was carried out for approximately 1 h using ethyl acetate, formic acid, acetic acid glacial, and water (100:12:12:26) as mobile phase in a closed glass tank. The chromatograms were examined under UV-A (365 nm, Fisher, Bioblock Scientific) and photographed to detect sinapate esters. Subsequently, separated phenylpropanoids were stained by spraying the plates with a 1% (w/v) diphenylboric acid 2-aminoethylester (DPBA; Roth) solution in 80% (v/v) methanol, followed by examination under UV-A (365 nm, Fisher, Bioblock Scientific) and documentation by photography. Several HPTLC-separated flavonol glycosides and sinapate esters were labelled according to a previous report that identified their structures by a successive combination of HPLC-PDA-ESI/MS (Stracke et al., 2007).

The same extraction method was used to prepare samples for absorption spectra measurements. After the centrifugation step, 100 µL of clear supernatant were transferred to a transparent 96-well plate and absorption spectra were measured using a TECAN Spark 10M Multimode Microplate Reader.

### Analysis of secondary metabolites by LC-MS/MS

Plant material was manually ground in a mortar under liquid nitrogen. Fifty-mg of powder were transferred into 1.5-mL tube, mixed with 5 volumes of 80% (v/v) methanol containing 0.1% (v/v) formic acid, and further homogenized using a TissueLyser II (Qiagen) and glass beads (diameter 2–3mm) for 3 min at 30 Hz. Samples were then centrifuged for 3 min at 12’000 *g*, clear supernatants were collected and transferred into HPLC vials for analysis. The targeted analysis of phenolic compounds was performed using ultrahigh performance liquid chromatography-photodiode array detection-high resolution mass spectrometry (UHPLC-PDA-HRMS). The system comprised an Acquity UPLC coupled to an eLambda PDA and a Synapt G2 QTOF (Waters) and was entirely controlled by MassLynx 4.1. The separation was performed on an Acquity UPLC CSH C18 column (100 x 2.1 mm, 1.7 µm, Waters) at a flow rate of 0.4 mL/min. A gradient of water + 0.05% (v/v) formic acid (phase A) and acetonitrile + 0.05% (v/v) formic acid (phase B) was applied as follows: 2–30% B in 6 min, 30–100% B in 3.5 min, hold at 100% (v/v) B for 1.5 min, reequilibration at 2% (v/v) B for 4 min. The PDA range was 190–600 nm and its acquisition frequency 5 Hz. The QTOF mass spectrometer was operated in electrospray negative ionization in full scan mode (scan range 50–1200 Da). Four flavonoids, four sinapoyl derivatives, coumaroyl-glucose, and feruloyl-glucose were profiled (Table S1-S3). Sinapoyl derivatives and other HCAs were quantified as sinapate equivalents and flavonoids as rutin equivalents, all using external calibrations (Glauser et al., 2012; Moreira et al., 2018). Data were processed in TargetLynx XS (Waters).

For the analysis of bands extracted from a HPTLC silica plate, the selected band was scraped off from the HPTLC plate and silica powder was resuspended in 80% (v/v) methanol and 0.1% (v/v) formic acid. Samples were vortexed 3 times for 10 sec and kept overnight at 4°C in the dark. Then samples were centrifuged 5 min at 12’000 *g* and supernatants were collected and transferred into HPLC vials for analysis. The untargeted analysis of HPTLC fractions was performed using a similar set-up with small modifications. The separation was performed on an Acquity UPLC HSS T3 column (100×2.1mm, 1.8 µm, Waters). The mass spectrometer was again operated in electrospray negative ionization but data-independent acquisition using the so-called MSe mode was used. The obtained chromatograms were visually inspected for differences and peaks of interest were identified based on their UV and MS/MS spectra. For the interpretation of MS/MS spectra, a combination of spectral matching using the RIKEN spectral database ReSpect and in-silico fragmentation using Sirius 5.5.1 including the CSI:FingerID and Canopus suite was used.

### Statistical analyses

Statistical analyses were performed using GraphPad Software Prism 9 software (San Diego, California). Statistical significance of the differences between means was determined using two-way analysis of variance (ANOVAs) followed by Tukey’s test for multiple comparisons, except for Figure 2A where log2 transformation of the values was performed before the ANOVA.

## ACCESSION NUMBERS

Sequence data from this work can be found in the Arabidopsis Genome Initiative or GenBank/EMBL databases under the following accession numbers: AT5G13930 (CHS/TT4), AT4G08920 (CRY1), AT1G04400 (CRY2), AT4G36220 (FAH1), AT5G11260 (HY5), AT3G17609 (HYH), AT3G62610 (MYB11), AT2G47460 (MYB12), AT5g49330 (MYB111), AT1G09570 (PHYA), AT2G18790 (PHYB), AT5G52250 (RUP1), AT5G23730 (RUP2), AT5G63860 (UVR8),

## AUTHOR CONTRIBUTIONS

M.L., E.D., and R.U. conceived and designed the research; M.L., F.A-O., and E.D. performed the experiments; G.G. performed and contributed metabolomics data; N.T. and R.P. generated and contributed genetic material; M.L., E.D., and R.U. analysed the data and wrote the paper. All authors reviewed and approved the submitted manuscript.

## ACKNOWLEDGEMENTS

We would like to thank Michel Goldschmidt-Clermont for helpful comments on the manuscript, Yamama Naciri , José Manuel Nunes and Tom Walker for advice on statistics, Henrik Johansson for kindly providing *hy5-215 hyh* mutant seeds, and the Nottingham Arabidopsis Stock Centre (NASC) for providing various other seed material. This work was supported by the University of Geneva and the Swiss National Science Foundation (grants 31003A_175774 and 310030_207716 to R.U.). R.P. was supported by an iGE3 PhD Salary Award.

## CONFLICT OF INTEREST STATEMENT

The authors declare that they have no conflict of interests.

## SUPPLEMENTAL DATA

**Supplemental Figure S1.**
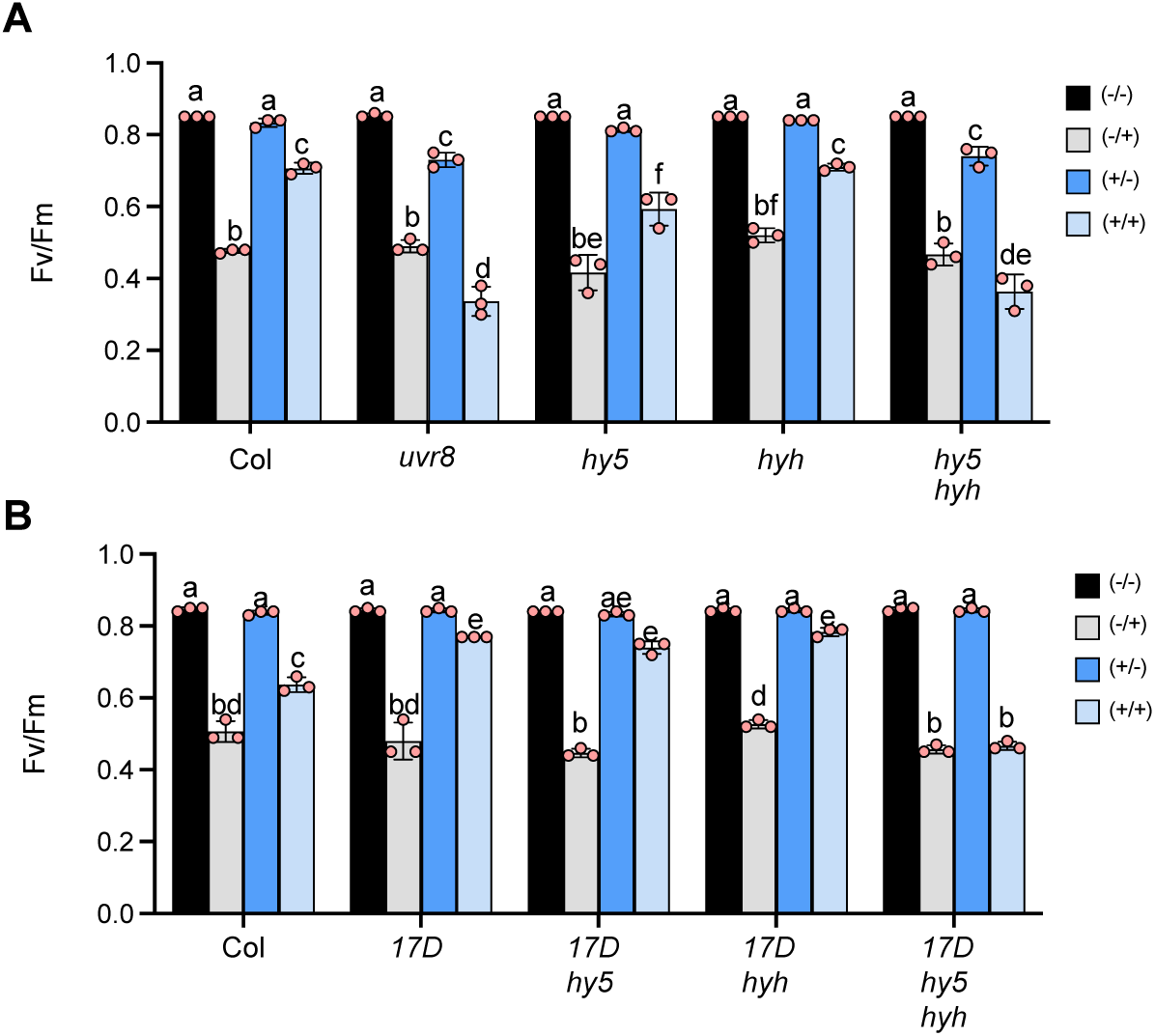
Enhanced photoprotection through UVR8-mediated UV-B acclimation depends on HY5 and HYH. **(A, B)** Maximum efficiency of PSII (Fv/Fm) in **(A)** wild type (Col), *uvr8-6* (*uvr8*), *hy5, hyh*, and *hy5 hyh*, and in **(B)** Col, *uvr8-17D* (*17D*), *uvr8-17D hy5* (*17D hy5*), *uvr8-17D hyh* (*17D hyh*), and *uvr8-17D hy5 hyh* (*17D hy5 hyh*) grown as described in Figure 1A. Values of independent experiments (orange points) and means ± SD are shown (*n* = 3). Shared letters indicate no statistically significant difference between the means (P > 0.05), as determined by two-way ANOVAs followed by Tukey’s test for multiple comparisons. -/-, not acclimated, not stressed; -/+, not acclimated, stressed; +/-, acclimated, not stressed; +/+, acclimated, stressed.

**Supplemental Figure S2.**
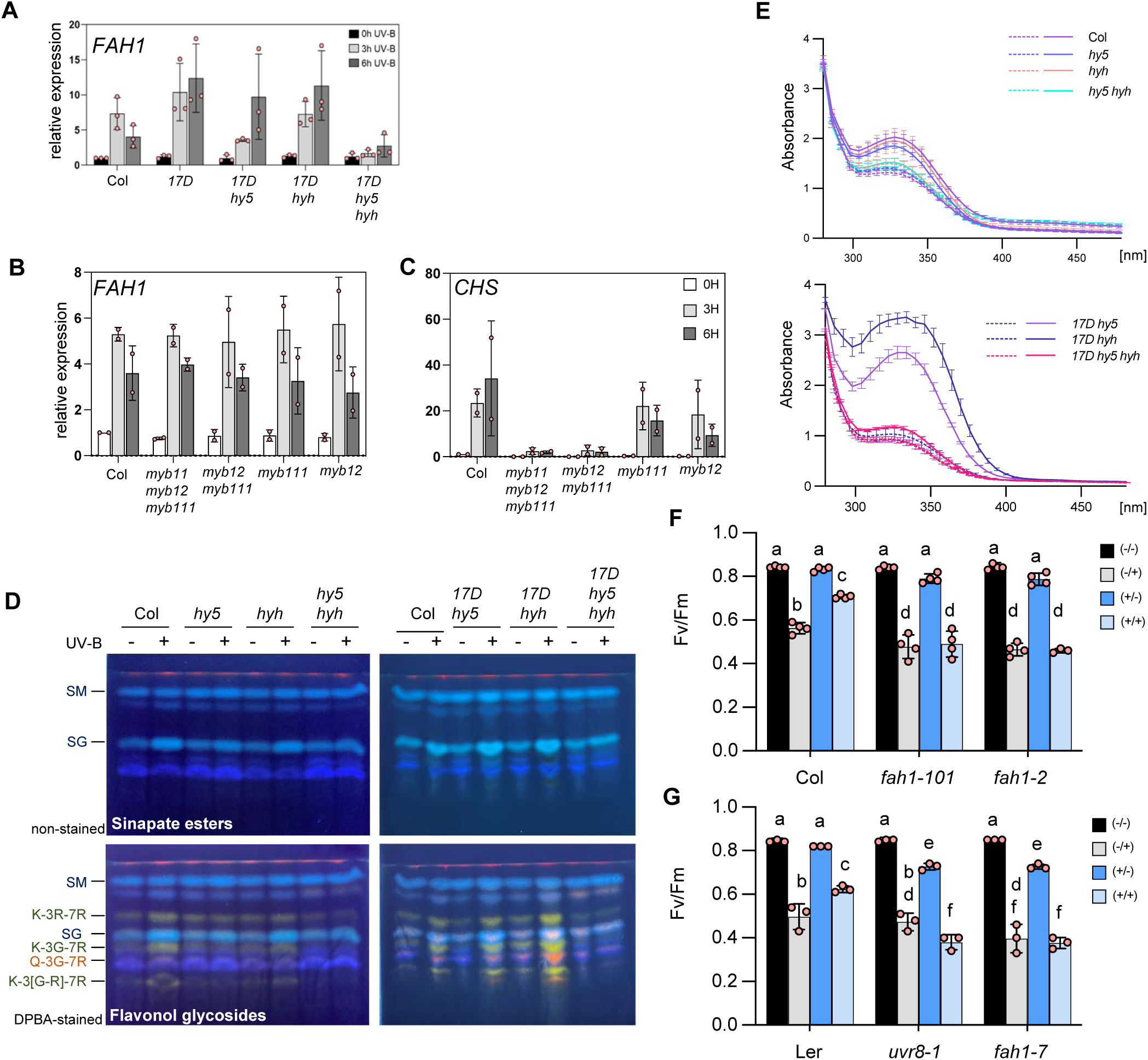
UVR8 signalling induction of *FAH1* expression and accumulation of sinapate esters depends on HY5 and HYH. **(A)** Quantitative RT–PCR of *FAH1* gene expression changes in response to 3h and 6h supplemental UV-B (0.08 mW cm^-2^) in 7-d-old light grown seedlings (20 µmol m^-2^ s^-1^) of upper panel : *uvr8-17D* (*17D*), *uvr8-17D hy5* (*17D hy5*), *uvr8-17D hyh* (*17D hyh*), and *uvr8-17D hy5 hyh* (*17D hy5 hyh*) compared to wild type (Col). Values of independent measurements (orange points) and means ± SDs are shown (*n* = 3) ; lower panel of *FAH1* and *CHS* gene expression in Col, *myb11 12 111, myb12 11, myb111, myb12.* Values of independent measurements (orange points) and means ± SDs are shown (*n* = 2). **(B)** HPTLC analysis of sinapate esters (*upper panels*) and flavonol glycosides (*lower panels*) levels in 10-d-old seedlings of wild type (Col), *hy5*, *hyh*, *hy5 hyh*, *uvr8-17D hy5* (*17D hy5*), *uvr8-17D hyh* (*17D hyh*), and *uvr8-17D hy5 hyh* (*17D hy5 hyh*) grown for 7 days in white light (20 µmol m^-2^ s^-1^) and exposed to supplemental UV-B (0.08 mW cm^-2^) for 3 days (UV-B: +) or not (UV-B: -). SM, sinapoyl malate; K-3R-7R, kaempferol-3-*O*-rhamnoside-7-*O*-rhamnoside; SG, sinapoyl glucose; K-3G-7R, kaempferol-3-*O*-glucoside-7-*O*-rhamnoside; Q-3G-7R, quercetin-3-*O*-glucoside-7-*O*-rhamnoside; K-3[G-R]-7R, kaempferol 3-*O*-[rhamnosyl-glucoside]-7-*O*-rhamnoside. **(C)** 280–480 nm absorption spectra of methanolic extracts from 10-d-old seedlings of wild type (Col), *hy5*, *hyh*, *hy5 hyh*, *uvr8-17D hy5* (*17D hy5*) , *uvr8-17D hyh* (*17D hyh*), and *uvr8-17D hy5 hyh* (*17D hy5 hyh*) grown for 7 days in white light (20 µmol m^-2^ s^-1^) and exposed to supplemental UV-B (0.08 mW cm^-2^) for 3 days (+, purple) or not (-, grey). Values of means ± SEM are shown (*n* = 3). **(D, E)** Fv/Fm measurements of 10-d-old seedlings of (D) wild type (Col), *fah1-101*, and *fah1-2*, and (E) wild type (Ler), *uvr8-1*, and *fah1-7* grown as described in Figure 1A. Values of independent experiments (orange points) and means ± SD are shown (*n* = 3 or 4). Shared letters indicate no statistically significant difference between the means (P > 0.05), as determined by two-way ANOVAs followed by Tukey’s test for multiple comparisons. -/-, not acclimated, not stressed; -/+, not acclimated, stressed; +/-, acclimated, not stressed; +/+, acclimated, stressed.

**Supplemental Figure S3.**
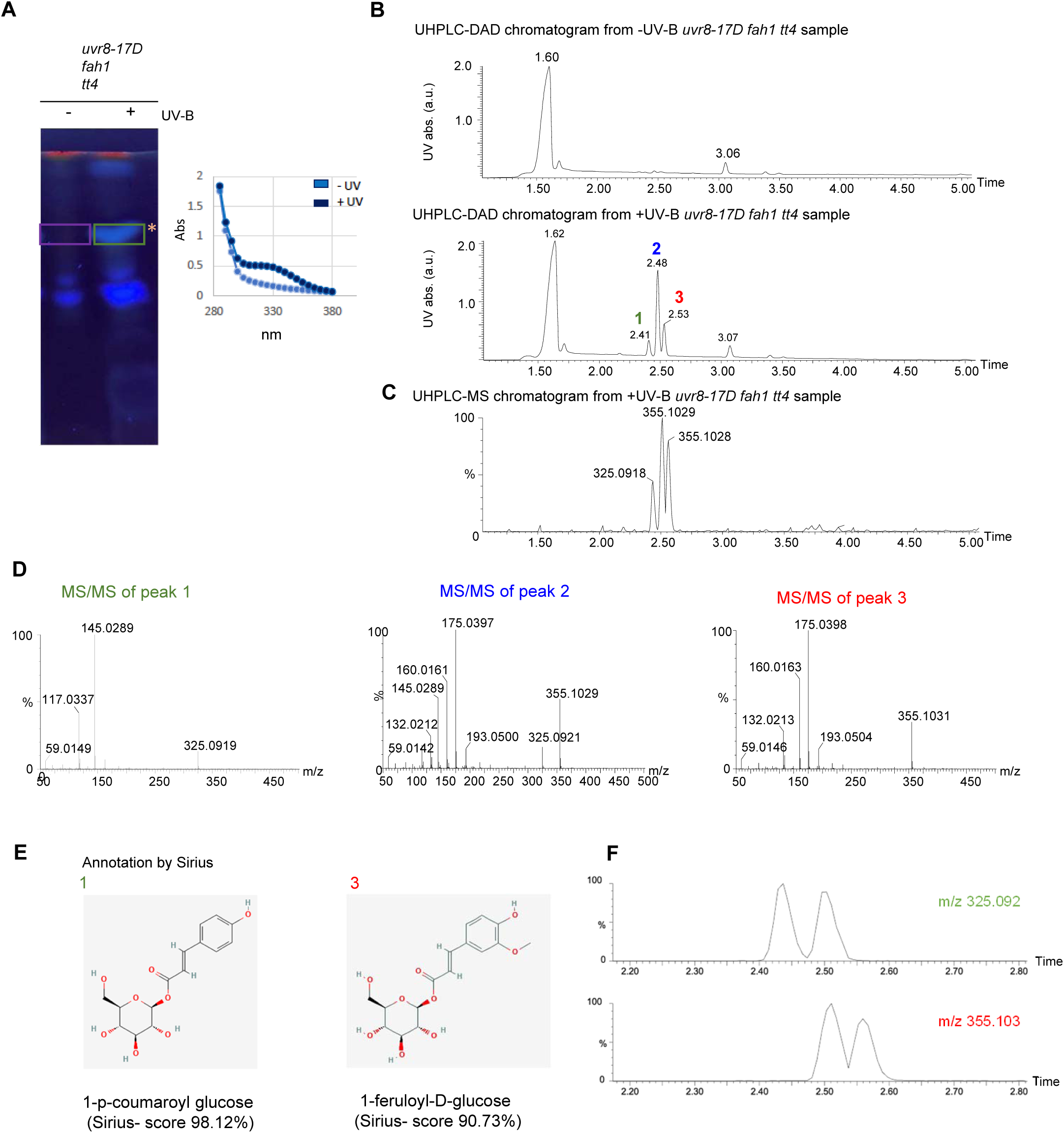
Enhanced UVR8 signalling promotes accumulation of coumaroyl glucose and feruloyl glucose. **(A)** (left panel) HPTLC of sinapate esters in 10-d-old *uvr8-17D fah1 tt4* seedlings grown for 7 days in white light (20 µmol m^-2^ s^-1^) and exposed to supplemental UV-B (0.08 mW cm^-2^) for 3 days (UV-B: +) or not (UV-B: -). The asterisk (*) indicates the specific metabolite(s) mainly detectable in the *fah1* mutant extracts (see Figure 3C, D); **(A)** (right panel) Absorption spectra between 280–480 nm of UV-absorbing compounds in methanolic extracts prepared from bands scraped off the silica plate at the indicated positions (-UV corresponds to purple rectangle on HPTLC, +UV corresponds to green rectangle). **(B)** UHPLC-DAD chromatograms of methanolic extracts from silica scraped off from the HPTLC plate region indicated by the purple rectangle in Supplemental Figure S3A (-UV-B) (*upper panel*) or from the region indicated by the green rectangle (+UV-B) (*lower panel*). a.u., arbitrary unit. 1 (green), 2 (blue), 3 (red) indicate peaks present in the + UV-B but not -UV-B extracts. **(C)** UHPLC-MS chromatogram of methanolic extracts from silica scraped off from the HPTLC plate region indicated by the green rectangle (Supplemental Figure S3A; +UV-B). **(D)** UHPLC-MS/MS spectra of peaks 1, 2 and 3 indicated in Supplemental Figure S3B. **(E)** Chemical structures of 1-p-coumaroyl glucose and 1-feruloyl-D-glucose, identified with the highest score by the program Sirius as the major compounds in peaks 1 and 3, respectively. **(F)** UHPLC-MS extracted ion chromatograms at m/z 325.092 and m/z 355.103 showing the presence of two isomers for both coumaroyl-glucose and feruloyl-glucose. Peak 2 contains a mixture both compounds detected in peaks 1 and 3.

**Supplemental Figure S4.**
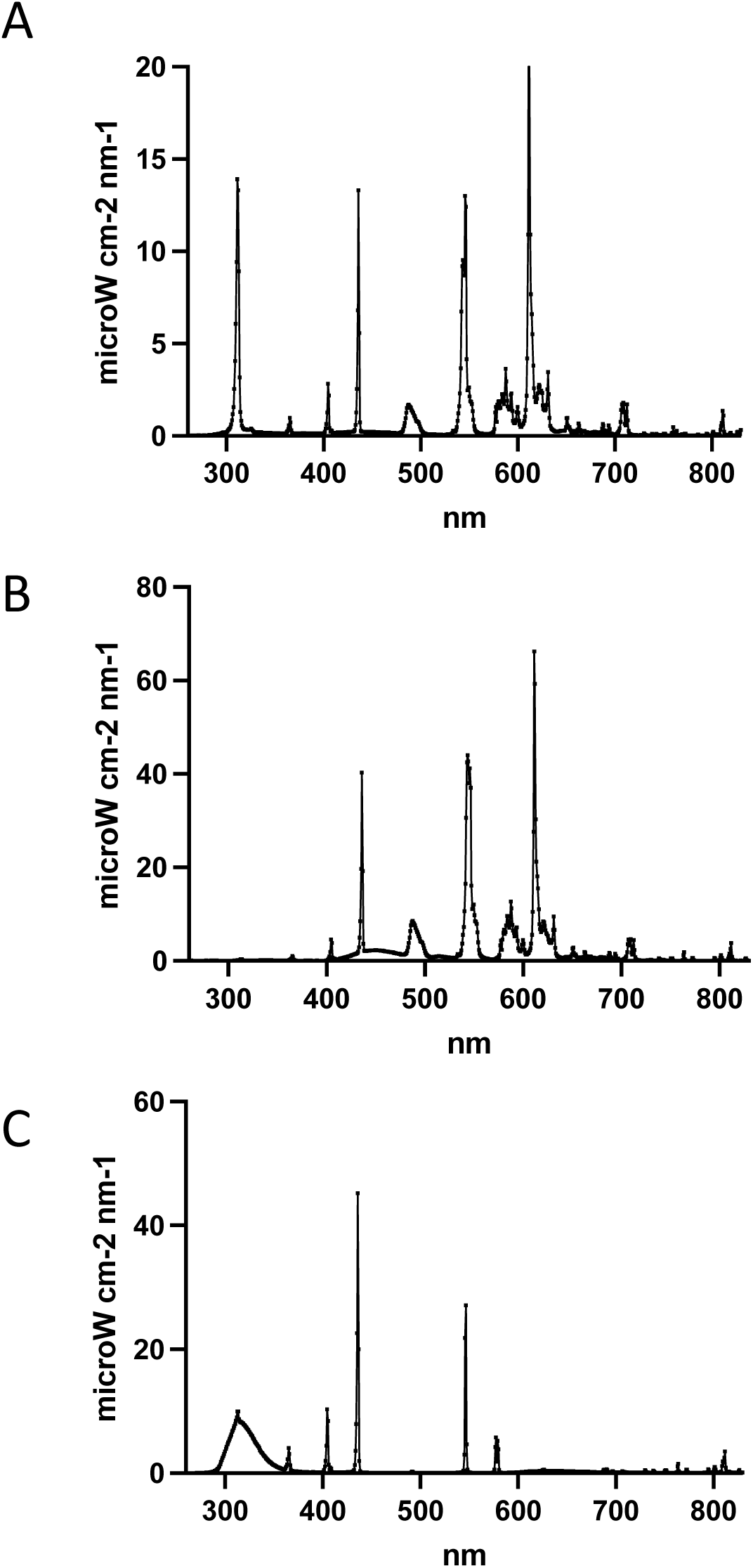
Spectra of the different light sources used in the study. **(A)** Spectra of the Osram L18W/30 white-light fluorescent tubes (20 μmol m^−2^ s^−1^), supplemented with Philips TL20W/01RS narrowband UV-B tubes (0.08 mW cm^−2^), **(B)** spectrum of Philips Alto II F17T8/TL841 white-light tubes (100 μmol m^−2^ s^−1^), **(C)** spectrum of the broadband UV-B tubes (Philips TL20W/12RS; 2.2 mW cm^−2^). **(A-C)** Spectral irradiance was measured in 0.8 nm intervals using an Ocean Optics QE65000 spectrometer.

**Supplemental Table S1.** Quantification of hydroxycinnamic acids (HCAs) and flavonol glycosides in 10-day-old seedlings grown in vitro (related to Figure 2).

| condition | genotype | Sinapoyl-glucose |  | Sinapoyl-malate |  | diSinapoyl-glucose |  | diSinapoyl-glucose isomer |  | K-3G-7R |  | Q-3G-7R |  | K-3R-7R |  | K-3[G-R]-7R |  |
| --- | --- | --- | --- | --- | --- | --- | --- | --- | --- | --- | --- | --- | --- | --- | --- | --- | --- |
|  |  | average | SD | average | SD | average | SD | average | SD | average | SD | average | SD | average | SD | average | SD |
| - UV | Col | 68.0 | 7.3 | 2.5 | 0.6 | 9.0 | 1.8 | 2.5 | 0.6 | 2.8 | 0.5 | 10.3 | 1.3 | 31.5 | 3.5 | 16.5 | 1.3 |
|  | *uvr8* | 68.3 | 5.2 | 4.3 | 0.5 | 8.0 | 0.8 | 3.0 | 0.0 | 3.3 | 0.5 | 12.3 | 1.0 | 37.3 | 2.2 | 19.5 | 2.5 |
|  | *rup1 rup2* | 64.8 | 8.2 | 4.3 | 0.5 | 9.3 | 2.1 | 3.5 | 0.6 | 4.5 | 0.6 | 10.0 | 1.8 | 32.5 | 4.2 | 17.8 | 2.4 |
|  | *uvr8-17D* | 62.3 | 8.5 | 5.8 | 0.5 | 8.8 | 0.5 | 2.5 | 0.6 | 5.5 | 0.6 | 12.8 | 1.3 | 34.8 | 3.1 | 17.5 | 1.7 |
|  | *hy5* | 90.3 | 9.6 | 2.3 | 0.5 | 14.3 | 1.9 | 6.0 | 0.8 | 0.8 | 0.5 | 6.8 | 1.0 | 23.0 | 2.2 | 17.5 | 3.1 |
|  | *hyh* | 66.5 | 5.9 | 4.0 | 0.8 | 8.0 | 1.4 | 2.0 | 1.4 | 4.0 | 1.2 | 12.5 | 3.1 | 39.0 | 7.9 | 15.3 | 2.8 |
|  | *hy5 hyh* | 101.8 | 7.9 | 4.5 | 0.6 | 10.0 | 1.2 | 4.3 | 1.0 | 1.3 | 1.3 | 8.8 | 1.3 | 24.8 | 4.1 | 12.3 | 1.7 |
|  | *fah1-101* | 19.0 | 14.9 | 0.3 | 0.5 | 1.0 | 1.2 | 0.5 | 0.6 | 2.8 | 0.5 | 6.8 | 1.0 | 26.3 | 4.4 | 10.0 | 0.8 |
|  | *tt4* | 61.0 | 3.6 | 4.5 | 0.6 | 3.8 | 0.5 | 1.5 | 0.6 | 0.8 | 1.0 | 0.8 | 1.0 | 5.3 | 3.4 | 2.3 | 2.2 |
| + UV | Col | 214.3 | 34.9 | 6.0 | 1.4 | 11.8 | 2.4 | 20.0 | 5.6 | 17.3 | 4.9 | 22.5 | 7.2 | 88.5 | 31.2 | 29.5 | 5.1 |
|  | *uvr8* | 73.0 | 9.1 | 4.8 | 0.5 | 7.0 | 0.8 | 2.0 | 0.8 | 3.5 | 0.6 | 13.5 | 1.7 | 39.0 | 3.7 | 21.0 | 1.4 |
|  | *rup1 rup2* | 556.0 | 14.6 | 7.8 | 0.5 | 21.0 | 3.8 | 56.3 | 2.1 | 128.5 | 13.8 | 104.8 | 18.5 | 367.8 | 35.8 | 151.3 | 17.2 |
|  | *uvr8-17D* | 349.5 | 23.1 | 9.3 | 1.3 | 16.3 | 1.5 | 29.3 | 4.1 | 58.8 | 7.8 | 44.5 | 11.4 | 188.0 | 20.3 | 69.3 | 5.5 |
|  | *hy5* | 130.3 | 37.2 | 4.0 | 1.4 | 9.5 | 2.6 | 8.0 | 2.4 | 1.3 | 0.5 | 7.3 | 1.0 | 24.8 | 4.3 | 17.0 | 2.2 |
|  | *hyh* | 218.5 | 50.4 | 7.5 | 2.4 | 10.5 | 2.5 | 17.0 | 5.9 | 18.8 | 5.1 | 29.8 | 6.2 | 91.3 | 19.6 | 32.5 | 7.2 |
|  | *hy5 hyh* | 99.8 | 6.8 | 4.3 | 0.5 | 8.0 | 2.2 | 3.0 | 0.0 | 0.3 | 0.5 | 6.5 | 1.3 | 19.0 | 2.2 | 10.3 | 1.7 |
|  | *fah1-101* | 0.0 | 0.0 | 0.0 | 0.0 | 0.0 | 0.0 | 0.0 | 0.0 | 16.8 | 1.3 | 19.8 | 2.1 | 79.5 | 7.0 | 24.5 | 1.3 |
|  | *tt4* | 228.0 | 35.3 | 11.5 | 3.7 | 6.8 | 0.5 | 22.5 | 7.9 | 0.0 | 0.0 | 0.0 | 0.0 | 0.0 | 0.0 | 0.0 | 0.0 |

Average in µg equivalent of sinapate /g FW for HCA esters, in µg equivalent of rutin /g FW for flavonol glycosides.
SD= Standard Deviation of the mean (n=4)
K-3G-7R, kaempferol-3-O-glucoside-7-O-rhamnoside; Q-3G-7R, quercetin-3-O-glucoside-7-O-rhamnoside; K-3R-7R, kaempferol-3-O-rhamnoside-7-O-rhamnoside; K-3[G-R]-7R, kaempferol 3-O-[rhamnosyl-glucoside]-7-O-rhamnoside

**Supplemental Table S2.** Quantification of hydroxycinnamic acids (HCAs) and flavonol glycosides in 10-day-old seedlings grown in vitro (related to Figure 3).

| condition | genotype | Sinapoyl-glucose |  | Sinapoyl-malate |  | diSinapoyl-glucose isomer 1 |  | diSinapoyl-glucose isomer 2 |  | Coumaroyl-glucose isomer 1 |  | Coumaroyl-glucose isomer 2 |  | Feruloyl -glucose isomer 1 |  | Feruloyl -glucose isomer 2 |  | K-3G-7R |  | Q-3G-7R |  | K-3R-7R |  | K-3[G-R]-7R |  |
| --- | --- | --- | --- | --- | --- | --- | --- | --- | --- | --- | --- | --- | --- | --- | --- | --- | --- | --- | --- | --- | --- | --- | --- | --- | --- |
|  |  | average | SD | average | SD | average | SD | average | SD | average | SD | average | SD | average | SD | average | SD | average | SD | average | SD | average | SD | average | SD |
| - UV | Col | 49.0 | 3.5 | nd |  | 6.0 | 1.6 | 0.0 | 0.0 | 0.0 | 0.0 | 0.0 | 0.0 | 0.0 | 0.0 | 0.0 | 0.0 | 3.5 | 1.0 | 7.0 | 1.2 | 29.5 | 3.4 | 17.0 | 2.6 |
|  | *uvr8* | 52.5 | 7.7 | nd |  | 7.5 | 1.0 | 0.0 | 0.0 | 0.0 | 0.0 | 0.0 | 0.0 | 0.0 | 0.0 | 0.0 | 0.0 | 7.5 | 1.9 | 14.5 | 3.4 | 52.0 | 8.3 | 26.0 | 3.7 |
|  | *rup1 rup2* | 60.0 | 5.9 | nd |  | 8.5 | 1.9 | 0.0 | 0.0 | 0.0 | 0.0 | 0.0 | 0.0 | 0.0 | 0.0 | 0.0 | 0.0 | 7.0 | 1.2 | 12.5 | 1.0 | 39.0 | 2.6 | 28.0 | 4.6 |
|  | *uvr8-17D* | 60.0 | 3.7 | nd |  | 10.0 | 0.0 | 0.0 | 0.0 | 0.0 | 0.0 | 0.0 | 0.0 | 0.0 | 0.0 | 0.0 | 0.0 | 12.5 | 1.0 | 17.0 | 2.6 | 60.5 | 2.5 | 33.5 | 1.9 |
|  | *fah1* | 0.5 | 1.0 | nd |  | 0.0 | 0.0 | 0.0 | 0.0 | 0.0 | 0.0 | 0.0 | 0.0 | 0.0 | 0.0 | 0.0 | 0.0 | 6.0 | 1.6 | 16.5 | 2.5 | 33.5 | 7.0 | 17.5 | 4.4 |
|  | *tt4* | 56.0 | 4.9 | nd |  | 4.0 | 0.0 | 0.0 | 0.0 | 0.0 | 0.0 | 0.0 | 0.0 | 0.0 | 0.0 | 0.0 | 0.0 | 0.0 | 0.0 | 0.0 | 0.0 | 0.0 | 0.0 | 0.0 | 0.0 |
|  | *uvr8-17D fah1* | 0.0 | 0.0 | nd |  | 0.0 | 0.0 | 0.0 | 0.0 | 0.0 | 0.0 | 0.0 | 0.0 | 0.0 | 0.0 | 0.0 | 0.0 | 7.5 | 1.0 | 12.0 | 1.6 | 31.5 | 2.5 | 23.0 | 1.2 |
|  | *uvr8-17D tt4* | 68.5 | 3.4 | nd |  | 4.5 | 1.0 | 0.0 | 0.0 | 0.0 | 0.0 | 0.0 | 0.0 | 0.0 | 0.0 | 0.0 | 0.0 | 0.0 | 0.0 | 0.0 | 0.0 | 0.0 | 0.0 | 0.0 | 0.0 |
|  | *uvr8-17D fah1 tt4* | 1.0 | 1.2 | nd |  | 0.0 | 0.0 | 0.0 | 0.0 | 0.0 | 0.0 | 0.0 | 0.0 | 0.0 | 0.0 | 0.0 | 0.0 | 0.0 | 0.0 | 0.0 | 0.0 | 0.5 | 1.0 | 0.0 | 0.0 |
| + UV | Col | 210.5 | 9.6 | nd |  | 8.0 | 0.0 | 10.5 | 3.8 | 0.0 | 0.0 | 0.0 | 0.0 | 0.0 | 0.0 | 0.0 | 0.0 | 26.5 | 4.7 | 35.0 | 7.7 | 113.0 | 21.7 | 42.0 | 2.8 |
|  | *uvr8* | 64.0 | 21.4 | nd |  | 3.5 | 2.5 | 0.0 | 0.0 | 0.0 | 0.0 | 0.0 | 0.0 | 0.0 | 0.0 | 0.0 | 0.0 | 8.5 | 1.0 | 16.0 | 1.6 | 54.0 | 8.2 | 32.5 | 10.9 |
|  | *rup1 rup2* | 605.0 | 44.0 | nd |  | 16.5 | 3.4 | 42.5 | 5.7 | 5.5 | 1.0 | 9.0 | 1.2 | 5.0 | 1.2 | 2.0 | 0.0 | 151.5 | 16.8 | 158.5 | 14.0 | 484.5 | 26.1 | 223.5 | 20.7 |
|  | *uvr8-17D* | 504.5 | 51.8 | nd |  | 15.0 | 1.2 | 33.0 | 4.2 | 2.0 | 0.0 | 3.5 | 1.9 | 1.5 | 1.0 | 0.0 | 0.0 | 105.0 | 10.6 | 103.0 | 14.3 | 336.5 | 24.6 | 136.0 | 13.9 |
|  | *fah1* | 0.0 | 0.0 | nd |  | 0.0 | 0.0 | 0.0 | 0.0 | 0.0 | 0.0 | 0.0 | 0.0 | 15.5 | 1.0 | 5.5 | 1.0 | 40.0 | 11.3 | 45.5 | 9.7 | 141.5 | 30.5 | 60.0 | 14.5 |
|  | *tt4* | 168.0 | 27.8 | nd |  | 6.5 | 1.9 | 9.0 | 3.8 | 0.5 | 1.0 | 0.5 | 1.0 | 0.0 | 0.0 | 0.0 | 0.0 | 0.0 | 0.0 | 0.0 | 0.0 | 0.0 | 0.0 | 0.0 | 0.0 |
|  | *uvr8-17D fah1* | 0.0 | 0.0 | nd |  | 0.0 | 0.0 | 0.0 | 0.0 | 5.0 | 1.2 | 8.0 | 2.8 | 63.0 | 12.3 | 21.5 | 3.0 | 100.0 | 27.9 | 70.5 | 18.0 | 332.5 | 74.2 | 144.5 | 41.5 |
|  | *uvr8-17D tt4* | 488.0 | 48.7 | nd |  | 13.5 | 4.4 | 46.0 | 6.3 | 14.0 | 3.7 | 18.5 | 3.0 | 9.5 | 1.9 | 3.5 | 1.0 | 0.0 | 0.0 | 0.0 | 0.0 | 0.0 | 0.0 | 0.0 | 0.0 |
|  | *uvr8-17D fah1 tt4* | 0.0 | 0.0 | nd |  | 0.0 | 0.0 | 0.0 | 0.0 | 36.5 | 4.7 | 48.0 | 5.4 | 240.5 | 25.3 | 89.5 | 5.5 | 0.0 | 0.0 | 0.0 | 0.0 | 0.0 | 0.0 | 0.0 | 0.0 |

Average in µg equivalent of sinapate /g FW for HCA esters, in µg equivalent of rutin /g FW for flavonol glycosides
SD= Standard Deviation of the mean (n=4)
K-3G-7R, kaempferol-3-O-glucoside-7-O-rhamnoside; Q-3G-7R, quercetin-3-O-glucoside-7-O-rhamnoside; K-3R-7R, kaempferol-3-O-rhamnoside-7-O-rhamnoside; K-3[G-R]-7R, kaempferol 3-O-[rhamnosyl-glucoside]-7-O-rhamnoside

**Supplemental Table S3.** Quantification of hydroxycinnamic acids (HCAs) and flavonol glycosides in soil-grown plants (related to Figure 4).

| condition | genotype | Sinapoyl-glucose |  | Sinapoyl-malate |  | diSinapoyl-glucose isomer 1 |  | diSinapoyl-glucose isomer 2 |  | Coumaroyl-glucose isomer 1 |  | Coumaroyl-glucose isomer 2 |  | Feruloyl -glucose isomer 1 |  | Feruloyl -glucose isomer 2 |  | K-3G-7R |  | Q-3G-7R |  | K-3R-7R |  | K-3[G-R]-7R |  |
| --- | --- | --- | --- | --- | --- | --- | --- | --- | --- | --- | --- | --- | --- | --- | --- | --- | --- | --- | --- | --- | --- | --- | --- | --- | --- |
|  |  | average | SD | average | SD | average | SD | average | SD | average | SD | average | SD | average | SD | average | SD | average | SD | average | SD | average | SD | average | SD |
| - UV | Col | 28.0 | 15.6 | 60.8 | 48.0 | 38.1 | 20.2 | 41.8 | 17.1 | 0.1 | 0.3 | 0.3 | 0.3 | 0.0 | 0.0 | 0.1 | 0.3 | 30.1 | 18.1 | 0.6 | 0.3 | 14.6 | 4.8 | 47.0 | 13.0 |
|  | *uvr8* | 19.8 | 4.4 | 50.1 | 4.2 | 19.6 | 21.6 | 23.9 | 19.2 | 0.0 | 0.0 | 0.3 | 0.3 | 0.0 | 0.0 | 0.0 | 0.0 | 19.1 | 3.8 | 0.5 | 0.0 | 11.5 | 2.0 | 39.6 | 3.3 |
|  | *uvr8-17D* | 62.3 | 11.6 | 48.8 | 11.3 | 33.5 | 26.0 | 29.9 | 15.6 | 0.1 | 0.3 | 0.4 | 0.3 | 0.0 | 0.0 | 0.0 | 0.0 | 88.4 | 29.7 | 49.0 | 27.8 | 123.8 | 57.4 | 110.5 | 31.7 |
|  | *rup1 rup2* | 77.8 | 11.8 | 58.1 | 9.6 | 39.3 | 34.0 | 35.3 | 20.0 | 0.4 | 0.3 | 0.5 | 0.0 | 0.0 | 0.0 | 0.0 | 0.0 | 155.3 | 17.9 | 112.3 | 14.8 | 236.1 | 21.0 | 171.3 | 10.5 |
|  | *tt4* | 39.3 | 4.6 | 63.0 | 7.6 | 28.6 | 27.8 | 31.7 | 23.0 | 0.1 | 0.3 | 0.5 | 0.0 | 0.1 | 0.3 | 0.0 | 0.0 | 0.9 | 0.5 | 0.9 | 0.5 | 2.0 | 1.1 | 1.5 | 0.7 |
|  | *fah1* | 0.9 | 0.5 | 0.5 | 0.4 | 0.6 | 0.2 | 0.4 | 0.2 | 0.0 | 0.0 | 0.0 | 0.0 | 0.8 | 0.3 | 1.0 | 0.0 | 29.0 | 4.1 | 1.9 | 0.8 | 21.0 | 3.1 | 60.6 | 5.6 |
|  | *fah1 tt4* | 0.4 | 0.3 | 0.3 | 0.3 | 0.3 | 0.1 | 0.2 | 0.1 | 0.0 | 0.0 | 0.0 | 0.0 | 0.4 | 0.3 | 1.0 | 0.0 | 0.5 | 0.0 | 0.3 | 0.3 | 0.6 | 0.3 | 1.1 | 0.6 |
|  | *uvr8-17D fah1* | 1.9 | 3.8 | 0.6 | 1.3 | 1.9 | 1.4 | 1.3 | 0.5 | 0.1 | 0.3 | 0.3 | 0.5 | 3.3 | 1.5 | 1.4 | 0.8 | 69.4 | 37.9 | 54.3 | 25.8 | 117.8 | 57.3 | 92.5 | 46.7 |
|  | *uvr8-17D tt4* | 91.8 | 27.4 | 53.8 | 22.1 | 48.7 | 31.9 | 39.1 | 14.7 | 1.5 | 1.1 | 1.9 | 1.2 | 0.9 | 0.5 | 0.3 | 0.3 | 1.1 | 0.9 | 0.8 | 0.9 | 2.0 | 1.8 | 1.5 | 1.4 |
|  | *uvr8-17D fah1 tt4* | 1.0 | 0.7 | 0.5 | 0.4 | 0.7 | 0.3 | 0.5 | 0.2 | 0.6 | 0.5 | 1.3 | 0.9 | 10.9 | 8.4 | 3.4 | 2.3 | 0.5 | 0.0 | 0.1 | 0.3 | 0.9 | 0.3 | 0.6 | 0.3 |
|  | *rup1 rup2 fah1* | 0.4 | 0.3 | 0.0 | 0.0 | 0.2 | 0.2 | 0.1 | 0.1 | 0.0 | 0.0 | 0.1 | 0.3 | 1.9 | 1.9 | 1.0 | 0.4 | 87.1 | 23.4 | 66.6 | 11.1 | 155.4 | 20.1 | 114.1 | 24.6 |
| + UV | Col | 99.0 | 10.7 | 89.8 | 16.3 | 54.0 | 46.9 | 51.7 | 30.1 | 27.0 | 4.4 | 31.4 | 4.1 | 6.6 | 0.6 | 2.5 | 0.4 | 97.4 | 10.9 | 92.0 | 12.3 | 155.6 | 13.0 | 111.1 | 7.7 |
|  | *uvr8* | 19.4 | 2.1 | 61.8 | 6.0 | 22.3 | 27.3 | 29.3 | 23.5 | 0.8 | 0.5 | 1.1 | 0.6 | 0.9 | 0.8 | 0.3 | 0.3 | 19.4 | 0.6 | 2.9 | 1.1 | 12.6 | 0.9 | 34.9 | 2.7 |
|  | *uvr8-17D* | 121.6 | 7.9 | 79.4 | 6.8 | 53.9 | 56.5 | 49.1 | 30.5 | 32.4 | 6.8 | 38.6 | 5.8 | 9.5 | 1.8 | 3.1 | 0.5 | 134.8 | 11.4 | 134.5 | 8.8 | 238.5 | 9.6 | 136.3 | 7.5 |
|  | *rup1 rup2* | 120.1 | 7.9 | 66.5 | 6.1 | 50.2 | 54.4 | 44.3 | 26.4 | 14.9 | 3.6 | 15.1 | 3.8 | 6.5 | 1.1 | 2.1 | 0.3 | 163.3 | 7.8 | 130.0 | 4.7 | 250.1 | 11.7 | 165.6 | 6.7 |
|  | *tt4* | 168.6 | 14.6 | 99.4 | 3.8 | 71.6 | 77.5 | 63.1 | 41.3 | 108.9 | 6.8 | 124.4 | 7.6 | 21.0 | 2.3 | 9.5 | 1.2 | 3.8 | 0.3 | 2.9 | 0.9 | 4.4 | 1.1 | 2.4 | 0.5 |
|  | *fah1* | 6.0 | 7.2 | 2.3 | 3.2 | 4.7 | 2.3 | 3.1 | 1.1 | 13.1 | 3.6 | 18.0 | 4.5 | 31.4 | 4.4 | 9.1 | 0.6 | 96.4 | 14.7 | 39.9 | 14.9 | 112.6 | 13.7 | 101.3 | 8.5 |
|  | *fah1 tt4* | 0.5 | 0.0 | 0.1 | 0.3 | 0.2 | 0.2 | 0.2 | 0.1 | 27.1 | 2.8 | 39.5 | 4.9 | 60.9 | 10.8 | 20.4 | 1.7 | 1.6 | 0.8 | 1.9 | 1.4 | 3.0 | 1.8 | 2.0 | 0.9 |
|  | *uvr8-17D fah1* | 1.5 | 2.0 | 0.5 | 0.4 | 1.1 | 0.8 | 0.7 | 0.3 | 38.9 | 3.6 | 43.5 | 3.9 | 74.0 | 3.5 | 16.8 | 1.4 | 144.9 | 8.2 | 130.3 | 7.0 | 227.1 | 8.6 | 130.5 | 3.8 |
|  | *uvr8-17D tt4* | 208.9 | 9.7 | 89.0 | 6.8 | 78.6 | 94.8 | 67.3 | 40.9 | 163.1 | 28.8 | 179.1 | 26.6 | 26.0 | 4.4 | 11.1 | 1.8 | 8.8 | 5.7 | 9.4 | 8.9 | 15.3 | 12.8 | 7.5 | 6.2 |
|  | *uvr8-17D fah1 tt4* | 2.6 | 1.9 | 1.3 | 0.9 | 1.7 | 0.8 | 1.1 | 0.4 | 137.1 | 3.7 | 159.6 | 5.0 | 170.0 | 6.1 | 53.8 | 3.2 | 2.5 | 2.1 | 3.9 | 3.6 | 5.9 | 4.7 | 3.0 | 2.5 |
|  | *rup1 rup2 fah1* | 2.4 | 1.9 | 2.5 | 3.7 | 2.6 | 0.8 | 2.4 | 1.2 | 9.6 | 1.7 | 10.6 | 1.4 | 52.1 | 4.9 | 11.1 | 1.2 | 152.0 | 2.9 | 114.9 | 2.9 | 218.4 | 7.9 | 145.3 | 3.2 |

Average in µg equivalent of sinapate /g FW for HCA esters, in µg equivalent of rutin /g FW for flavonoids
SD= Standard Deviation of the mean (n=4)
K-3G-7R, kaempferol-3-O-glucoside-7-O-rhamnoside; Q-3G-7R, quercetin-3-O-glucoside-7-O-rhamnoside; K-3R-7R, kaempferol-3-O-rhamnoside-7-O-rhamnoside; K-3[G-R]-7R, kaempferol 3-O-[rhamnosyl-glucoside]-7-O-rhamnoside

**Table S4:**
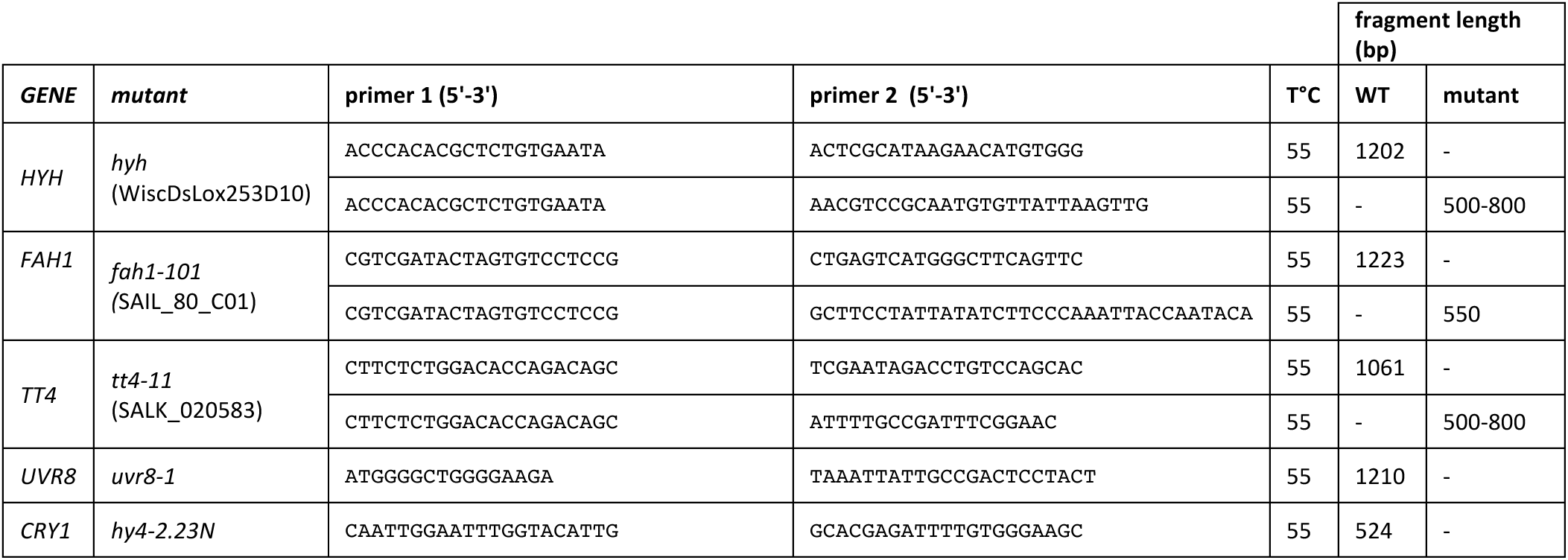
Genotyping information

## Notes

### Competing Interest Statement

The authors have declared no competing interest.

